# Increased Cognitive Load Expands, Stabilizes, and Aligns Visual Representation Across the Human Brain

**DOI:** 10.64898/2026.11.05.764028

**Authors:** Shijie Qu, Marvin M. Chun, Yaoda Xu

**Affiliations:** Department of Psychology, Yale University, New Haven, CT, USA; Wu Tsai Institute, Yale University, New Haven, CT, USA; Department of Neuroscience, Yale School of Medicine, New Haven, CT, USA

**Keywords:** Cognitive Load, Visual Representations, Multivariate Decoding, Neural Variance, Brain Networks

## Abstract

Cognitive load refers to the mental effort required to perform a task. While previous research has extensively examined its detrimental effects on neural and behavioral performance, much less is known about how the brain actively adapts to heightened demands. Across two fMRI experiments involving visual n-back tasks, we identified several fundamental properties of this adaptive process. First, higher load enhanced and expanded visual representations beyond localized visual regions, leading to widespread visual coding. At the global scale, heightened load promoted cross-regional coordination by increasing representational alignment and strengthening information encoded in interregional functional connectivity patterns. Beyond facilitating stronger visual representation, increased load also reduced neural pattern variability, which led to more stable information coding. Finally, load was encoded independently of other task-relevant information as a separate brain state, supporting its overarching role in guiding visual processing. Together, these findings revealed that the brain adapts to heightened cognitive demands by globally reorganizing neural processing to support flexible cognition.

**Highlights:**

- The brain employs multiple neural mechanisms to adapt to heightened cognitive demands.
- Increased cognitive load strengthens and expands visual representations beyond visual regions.
- Higher cognitive load enhances cross-regional coordination through increased representational alignment and stronger information coding in interregional connectivity patterns.
- Neural pattern variability decreases under greater load, promoting more stable representations
- Load is encoded as a distributed brain state, with minimal interaction with other variables

## Introduction

Every task we perform involves mental effort, a concept known as cognitive load (Chun and Most, 2026; Sweller, 1988). Cognitive load has been widely studied and is linked to robust activity in the higher-level cognitive control regions (Braver et al., 1997; Carlson et al., 1998; Duncan, 2010; Jansma et al., 2000; Jonides et al., 1997). Prior studies have largely focused on the impairment under high cognitive load when human performance is pushed to its limit. What is largely underexplored, however, is how the human brain dynamically copes with such demand, still manages to extract and represent task-relevant information, and succeeds in many circumstances. Understanding this adaptive capacity is essential to explaining how we flexibly and successfully handle the varying demands of everyday tasks.

Cognitive load has traditionally been associated with impaired cognition at both behavioral and neural levels. Higher cognitive load, typically induced by increased working memory demands in tasks such as the n-back task (Kirchner, 1958), leads to lower response accuracy and weaker suppression of task-irrelevant distractors (de Fockert et al., 2001; Lavie & De Fockert, 2005; Park et al., 2007; Yi et al., 2004). Beyond diminished neural suppression of distractors (Kelley & Lavie, 2011), higher cognitive load has also been associated with inattentional blindness (Todd et al., 2005) and lower precision of working memory contents (Chopurian et al., 2026; Emrich et al., 2013; Sprague et al., 2014). Together, these findings have contributed to a prevailing view of cognitive load as a constraint on visual cognition, whereby increasing task demands exhaust limited cognitive resources and compromise information processing.

However, rather than focusing on task failures, which are bound to happen when increased cognitive loads are taxing the limits of human cognition, it is more critical to understand how the brain sustains performance by adapting to demanding circumstances. One effective strategy would be to deploy additional resources and reorganize how task-relevant visual information is represented. Previous studies have offered some preliminary evidence for these adaptive processes. For instance, higher cognitive demand activates a broader network of brain areas with more efficient functional connections among them (Kitzbichler et al., 2011; Shine et al., 2016). Heightened cognitive effort has also been linked to stronger interactions between prefrontal and visual cortices, through which effort-related signals modulate the salience of visual information coding (Master et al., 2024). Our group has also observed enhanced decoding of visual information under higher cognitive load across predefined regions in the larger occipitotemporal cortex (OTC) and posterior parietal cortex (PPC; Xu & Chun, 2025).

Although these findings provide initial evidence that the brain may adaptively respond to heightened cognitive demands, our understanding of how cognitive load may actively shape information representation in the human brain remains limited. Previous studies have largely focused on specific effects within a restricted set of brain regions, leaving open the broader question of whether and how cognitive load induces more widespread and diverse changes in information representations across the cortex. Addressing this gap is critical for understanding not merely which brain regions are recruited under greater demand, but how the whole brain adapts and reorganizes its representations of task-relevant information to support cognition under challenging scenarios.

In this study, we employed a set of analytical approaches to systematically characterize the multiple ways in which cognitive load reshapes visual representations across the entire cortex. In particular, we analyzed four properties of how increased cognitive load reconfigures neural coding. First, we investigated the general impact of load on the strength and spatial extent of visual representations across the brain. We hypothesized that, under heightened cognitive demands, the brain would not only strengthen visual representations in regions previously implicated in load-dependent modulation, such as PPC and OTC (Xu & Chun, 2025), but also recruit a broader set of regions beyond the canonical ones, as has been observed for visual representations in naturalistic settings (Çukur et al., 2013; Huth et al., 2012).

Second, as suggested by earlier work (Kitzbichler et al., 2011; Shine et al., 2016), cognitive load can also influence how different parts of the brain synchronize with each other. We predicted that, in addition to magnitude changes in functional connectivity, cognitive load could further induce more profound representational changes across the brain in two ways. One involves a global reorganization of visual representational structures across different brain regions, with high load promoting greater alignment and integration rather than further functional differentiation (e.g., J. R. Cohen & D’Esposito, 2016). As a novel prediction, we hypothesize that increased integration will be accompanied by greater similarity in visual representations across cortical regions. Another way that cognitive load can affect visual representation is by changing the information encoded in functional connectivity patterns between brain regions. Moving beyond representations within individual regions, we explored whether the fine-grained patterns of functional connectivity between regions could themselves carry task-relevant representational information, and whether the strength of such representations could also be modulated by load. To test this possibility, we introduced a new method, functional connectivity pattern analysis (FCPA), to characterize inter-regional multivariate representations as an additional representational scheme and examine how they are modulated by cognitive load.

Third, besides pattern decoding accuracy, we examined another important property of neural representations: the variability of population coding. Previous electrophysiological studies have extensively investigated neural variability, showing that neural responses become less variable under certain task conditions and that such reductions in variability may facilitate response preparation and sustain performance in demanding situations (Churchland et al., 2006, 2010; M. R. Cohen & Maunsell, 2009; Mitchell et al., 2009). Despite these findings, neural variability has received relatively little attention in fMRI research (He, 2013; Ito et al., 2020). More importantly, while multivariate fMRI activity patterns have been extensively studied for the information they encode, remarkably little is known about how the variability of these patterns changes with cognitive demands. In this study, we extended the investigation of neural variability from single-neuron responses to population-level activity patterns measured with fMRI. We hypothesized that, in response to increased cognitive load, the brain would adopt more stable population-level representations to scaffold efficient information processing and sustain performance under heightened external task demands.

Finally, we examined how cognitive load itself, as an overarching task signal, is encoded across the brain during visual cognitive processing. Previous research has largely focused on increased but relatively localized activation in frontoparietal regions under higher cognitive load (Braver et al., 1997, 1997; Carlson et al., 1998; Duncan, 2010; Jansma et al., 2000). Drawing on recent evidence that load-related signals are present throughout the visual hierarchy (Adam et al., 2025) as well as findings demonstrating the widespread influence of general arousal signals across the brain (Raut et al., 2025), we hypothesized that cognitive load signals may be distributed across the majority of the cortex and could be decoded from multivariate activity patterns. We further predicted that as a modulatory signal, cognitive load would be encoded largely independently of other task-relevant information such as object and attentional state, reflecting a separable neural state rather than being represented solely through its modulatory effects on other neural representations.

To study these four properties of cognitive load on neural representations, we conducted two fMRI studies, one re-analyzed from Xu & Chun (2025), and the other newly collected. In both studies, participants completed 1-back (low load) or 2-back (high load) visual working memory tasks (Chun and Most, 2026; Sweller, 1988). Multivariate voxel and connectivity analyses were employed to systematically quantify each effect. Together, these findings offered an alternative view to the established resource-depletion account of cognitive load: as load increases, task-relevant visual information becomes more broadly represented and coordinated across the brain, giving rise to sharpened information readout under heightened cognitive demands.

## Results

To investigate neural representations of cognitive load on visual cognition, we first expanded the analyses of the fMRI data from a previous study (Xu & Chun, 2025; hereafter referred to as Experiment 1) where 12 participants performed an n-back task (Figure 1A, Supplementary Figure 1A, B), attending to objects or color under low (1-back) or high load (2-back). Each n-back block consisted of a sequence of ten images drawn from one of four object categories. To better characterize the fine-grained object representational structures in different brain regions across high or low load, in a second experiment (Experiment 2) designed for this study with 12 participants, we increased the number of object categories to eight (Supplementary Figure 2A). As in Experiment 1, we manipulated cognitive load using separate runs of 1-back and 2-back tasks, but we did not manipulate attention, and participants always performed the n-back task on the objects; thus, we presented the grayscale object images directly without any color dot overlay.

**Figure 1.**
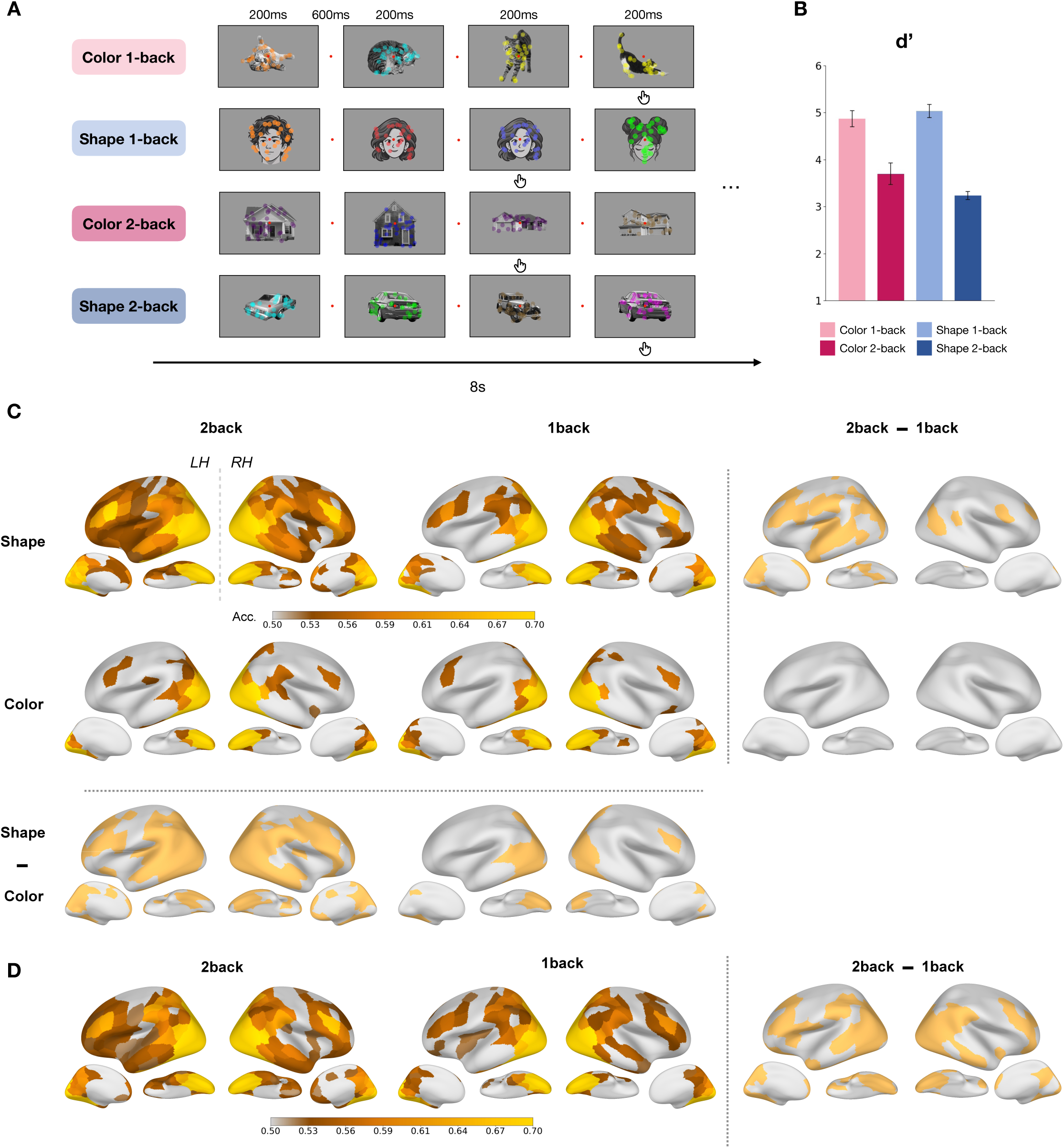
Task Design and Pattern Representations of Object Category Information. **A.** Sample *n*-back trial blocks for each of the four task conditions. Within each block, a sequence of images from a single object category was presented. Each stimulus consisted of a grayscale object overlaid with colored dots. The four object categories were cats, cars, faces, and houses. The cartoon faces shown here were generated using ChatGPT for illustration; the actual experiment used photographs of real human faces. The click icon indicates the target stimuli for which participants were required to respond when detecting an n-back repeat. **B**: D’ behavioral performance under each of the four task conditions (n=12). **C**: Whole-brain decoding maps for object category in the left and right hemispheres, shown separately across task conditions. Contrast maps on the right and bottom highlight regions with significant decoding differences across load and attention levels. Decoding accuracy in each region was tested against chance level, 0.5, using one-sample t-tests followed by FDR correction across all ROIs in each hemisphere. Contrast maps were generated using one-sided t-tests followed by FDR correction across all ROIs in each hemisphere. Regions that did not survive correction are shown in light gray. **D**: Whole-brain decoding maps for object category for Experiment 2.

Overall, participants performed better under low load than high load. In Experiment 1, Behavioral accuracy (*d’*) across the four task conditions (Figure 1B) showed a significant main effect of load (F = 232.971, p < 0.001), as well as a significant load × attention interaction effect (F = 7.904, p = .017). Similar results were obtained when *a’*, instead of d*’*, was used (Supplementary Figure 1C; F’s > 15.60, p’s < .002 for the main effect and the interaction). The performance drop with increased load was more pronounced when attention was directed to shape than to color. In Experiment 2, d’ and a’ are also significantly higher for 1-back than 2-back conditions (Supplementary Figure 2B; t’s > 7.86, p’s < 0.001).

### Cognitive Load Promotes Heightened and Distributed Visual Representations

We first sought to address the critical question of whether and how cognitive load modulates object representations across the brain. Using Experiment 1 data, we examined the neural coding of object category information using multivariate pattern analysis (MVPA), which identifies brain regions whose voxel activity patterns can reliably distinguish between different object categories.

Under different load conditions, we found that the spatial extent of object representations across the cortical surface changed markedly (Figure 1C): During the 1-back condition, when attention is deployed to the shape of the object (shape 1-back), object category was decodable in a relatively restricted set of PPC and OTC regions, along with a few lPFC regions (43 left, 54 right; based on the Schaefer-200 atlas, with 100 parcels per hemisphere). Strikingly, under high cognitive load (shape 2-back), object category decoding became most widespread across the cortex, extending across large portions of the lateral and ventral surfaces, as well as medial parietal and occipital cortices (90 left, 76 right; 109.3% and 40.7% increases, respectively). A direct comparison between the two conditions further confirmed that higher load was associated with stronger object representations across lateral prefrontal, parietal, and occipitotemporal regions.

Such a load-enhancing effect, however, was not present when attention was directed to the overlaid dots sharing the same spatial envelope rather than the object. During the color 1-back condition, object category decoding was largely confined to OTC, with only limited involvement of lPFC and PPC (27 left hemisphere ROIs, 30 right hemisphere ROIs). Increasing task load (color 2-back) resulted in only a modest increase in the number of significant ROIs (28 left, 32 right; 3.7% and 6.7% increases, respectively), and did not yield significantly higher decoding accuracy in any ROI when contrasting directly between the two conditions (all p’s > 0.05, FDR-corrected). These findings suggest that the facilitating effect of load on object representations depends on the attended feature, and is largely absent when attention is not directed to the relevant object feature.

Consistent with these observations, repeated-measures ANOVA showed that object category decoding was modulated by load, attention, and also their interaction across different cortical regions (Supplement Figure 1D). Effects of load were observed in a subset of PPC and lPFC regions in the left hemisphere and a subset of OTC regions in the right hemisphere (p’s < 0.05, FDR-corrected). Effects of attention were seen in the OTC, PPC, and lPFC that span the temporal, parietal, dorsal attention, and visual networks (p’s < 0.05, FDR-corrected). Finally, an interaction between load and attention emerged in the lPFC as well as the medial occipital and parietal cortex in both hemispheres (p’s < 0.05, FDR-corrected). These interaction effects confirmed that the load-dependent enhancement of object representations was more pronounced when attention was directed to the objects themselves rather than to the overlaid color information. Overall, these results demonstrated that the specific set of regions carrying object information can extend far beyond the canonical visual cortex under higher load, indicating the crucial role of load in facilitating a more distributed representation of visual information.

Besides object representation, increased load also makes attentional state more decodable across wider cortical areas. Under low load, attention decoding was significant across lateral frontal, parietal, and occipital regions, as well as the ventral occipital cortex (Supplementary Figure 3A; 45 left, 38 right; p’s < 0.05, FDR-corrected). Under high load, just as with object decoding, attentional state became decodable across a broader set of regions in OTC, lPFC, and medial parietal and occipital cortices (83 left, 68 right; 84.4% and 78.9% increases, respectively). A separate paired t-test further confirmed significantly increased attentional state decoding under 2-back across multiple cortical regions (28 left, 9 right; p’s < 0.05, FDR-corrected). Together, these results support the account that higher load also produces more distinct patterns for higher-level attentional states, in addition to the perceived object information.

Experiment 2 replicated the key finding that cognitive load enhances the spatial spread of attended object representations (Figure 1D, Supplementary Figure 2C). We increased the object categories from four to eight to rule out the possibility that the enhanced object decoding under higher load was driven simply by behavioral performance differences across object categories (Supplementary Figure 4A). Specifically, even when the analysis was restricted to a subset of object categories with matched behavioral performance, the more distributed object representations under high load remained evident (Supplementary Figure 4B), suggesting that the effect reflects a genuine enhancement of object representations rather than higher-level task differences.

### Increased Load Aligns Whole-Brain Fine-grained Object Representational Structure

While higher load enhances the strength of object representations in individual brain regions, it remains unknown how it affects information representation at the global level, whether it promotes representational differentiation (Christophel et al., 2017; D’Esposito & Postle, 2015), or instead drives representational convergence across brain regions (J. R. Cohen & D’Esposito, 2016). To examine these possibilities, we performed representational similarity analysis (RSA, Kriegeskorte et al., 2008) on Experiment 2 data, which included a larger set of object categories with participants performing a visual object n-back task without explicit attention manipulation.

With the larger number of object categories in Experiment 2, we first constructed representational dissimilarity matrices (RDM) across all object categories and load conditions for different ROIs and networks. The RDMs were obtained from the decoding accuracy between each condition pair, as a proxy for dissimilarity between their corresponding neural activity patterns. We then visualized these RDMs in a two-dimensional space for a few selected regions in lPFC, PPC, and OTC (Supplementary Figure 5A, B). As expected, these regions exhibited clustering primarily along their canonical representational dimensions, with lPFC and PPC showing stronger separation by load and OTC showing stronger separation by object category. Nevertheless, the relative arrangements of object categories also exhibited subtle changes between low and high load, indicating that increased cognitive load can reshape the fine-grained object representational structures.

To directly characterize load-induced representational changes on object representations, we derived an object RDM for each ROI under each load condition. We then calculated pairwise correlations between these RDMs to construct a larger ROI × Load RDM that captures the representational relationships across brain regions and load conditions. The resulting RDM was then projected into a two-dimensional space using MDS to visualize how object representations shifted across load conditions (Figure 2A). As ROIs belonging to the same functional network tended to cluster together in representational space (Supplementary Figure 6A, B), we streamlined this analysis to the network level by averaging object RDMs across ROIs within each network and repeating the same procedure (see Figure 2B for representative visualizations of network-level RDMs). This allowed us to more concisely characterize large-scale changes in the organization of object representations across the cortex.

**Figure 2.**
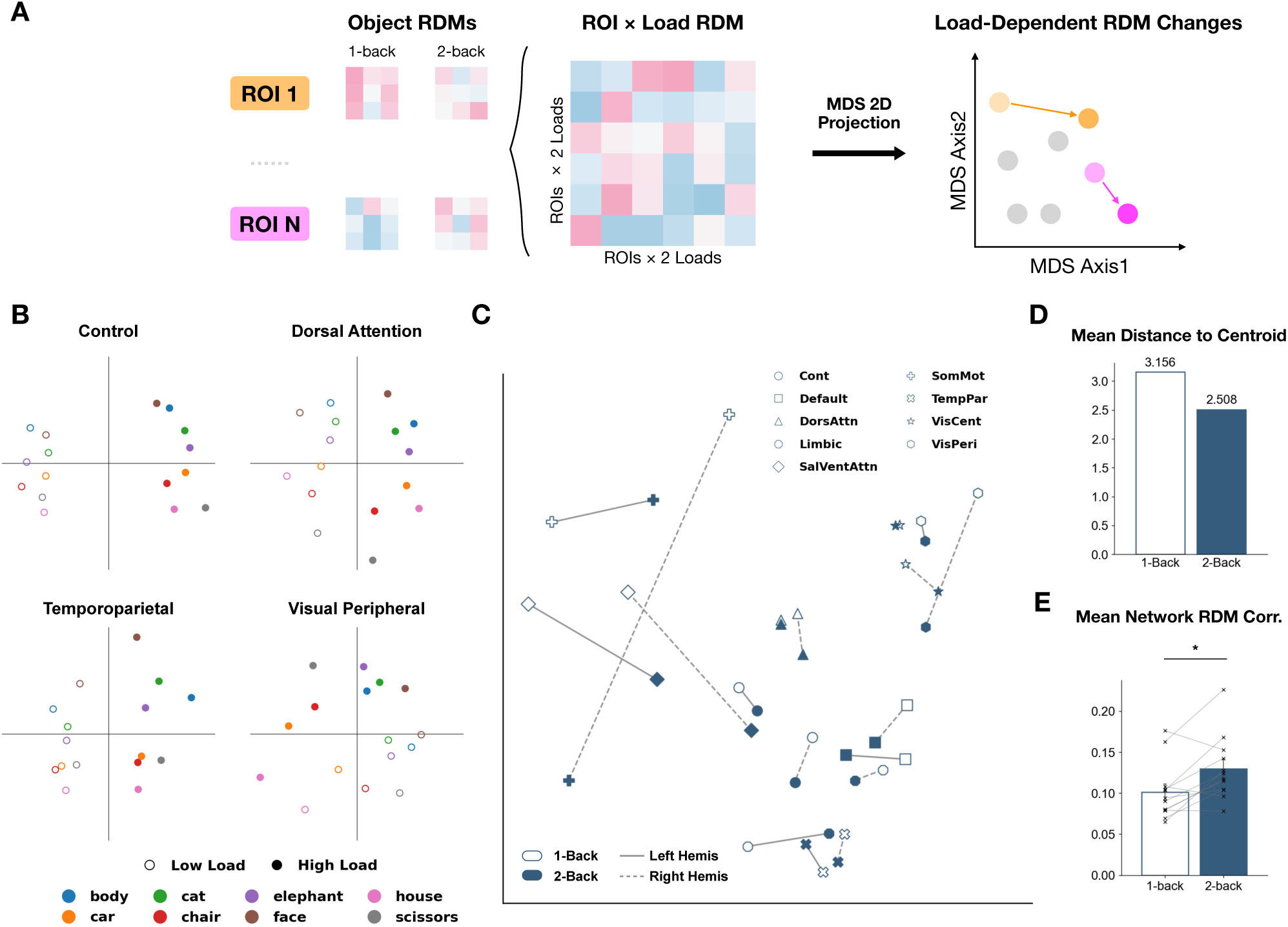
Load Aligns Cortex-Wide Object Representational Space. **A**: Schematic of the object RSA analysis. Object RDMs were constructed for each ROI (or network) under the two load conditions and used to derive a combined ROI (or network) × load RDM. The resulting RDM was then projected into a two-dimensional space using multidimensional scaling (MDS) to visualize load-dependent changes in object representational structures across ROIs (or networks). **B**: Sample MDS visualization of neural representations for each object category × load condition within selected functional networks in the left hemisphere. Dot color indicates object category, while marker fill (filled or hollow) indicates load condition. **C**: Network-level RDM shifts across cognitive loads. For each network, object-category RDMs were averaged across constituent ROIs, and MDS was performed on the resulting network-level RDMs to visualize their relationships in a two-dimensional space. Each marker represents one network, with hollow and solid markers indicating its coordinates under 1-back and 2-back conditions, respectively. Arrows connect the two load conditions to illustrate shifts in network-level representational structure. Left and right hemispheres were analyzed separately, yielding two arrows per network. Solid and dashed lines indicate the left and right hemispheres, respectively. **D**: Average distance to the centroid across networks under each load condition (based on panel C). To quantify alignment, we computed the Euclidean distance between each network’s MDS coordinate and the centroid (mean coordinate across all networks) within a given load condition. The bar plot displays average distance across networks for each load. **E**: Average RDM correlation across networks under each load. Each point represents one participant’s mean RDM correlation across networks, with values from the same participant across the two load conditions connected by a line. Only regions with significant average object category decoding accuracy were included. (Paired t-test: t = −3.388, p = 0.0061)

After projecting the network object RDM under each load simultaneously onto a 2D space (which captured the majority of the variance; Supplementary Figure 6I, J), we found that higher load promotes, rather than suppresses, object representational alignment across the brain (Figure 2C). This increased alignment was evidenced by a decrease in the mean distance of individual networks to the centroid of all networks (Figure 2D, Supplementary Figure 6C), as well as an increase in mean network RDM correlations (Figure 2E; t = −3.388, p = 0.006) under high load. Importantly, this effect was not driven solely by regions that only became significant in object decoding under high load, as excluding these regions yielded the same pattern of results (Supplementary Figure 6D-H). Together, these findings provide key evidence to arbitrate between the two opposing accounts of representational organization. Our results indicate that increasing cognitive load favors representational convergence over differentiation, leading distributed cortical systems to adopt a more coherent object representational code in support of a shared computational goal.

Meanwhile, this increased alignment does not eliminate differences in representational structure across networks, as shown by the reduced but still present separation between network coordinates (Figure 2C, Supplementary Figure 6E). These remaining differences indicate that different brain regions still retain their distinctive functions, as seen under less demanding conditions, while simultaneously becoming more coordinated to meet increased task demands.

### Load Effect on Visual Representations Through Inter-regional Connectivity Patterns

In parallel with globally transforming object representational structure, cognitive load may also meaningfully alter how different brain regions communicate with one another, thereby enabling better readout of object information under higher load. To test this hypothesis, we developed a novel approach, which we term functional connectivity pattern analysis (FCPA), that closely parallels MVPA. Using FCPA, we examined whether the multivariate voxel-to-voxel connectivity patterns between pairs of ROIs or networks can also reliably decode the three variables of interest (Figure 3A). Unlike other multivariate functional connectivity-based methods (Coutanche & Thompson-Schill, 2013; Nieto-Castanon, 2022), our approach does not transform connectivity into a latent or secondary feature space. Rather, it directly leverages patterns of pairwise correlation values, preserving voxel-level detail that has proven useful for characterizing representations in MVPA.

**Figure 3.**
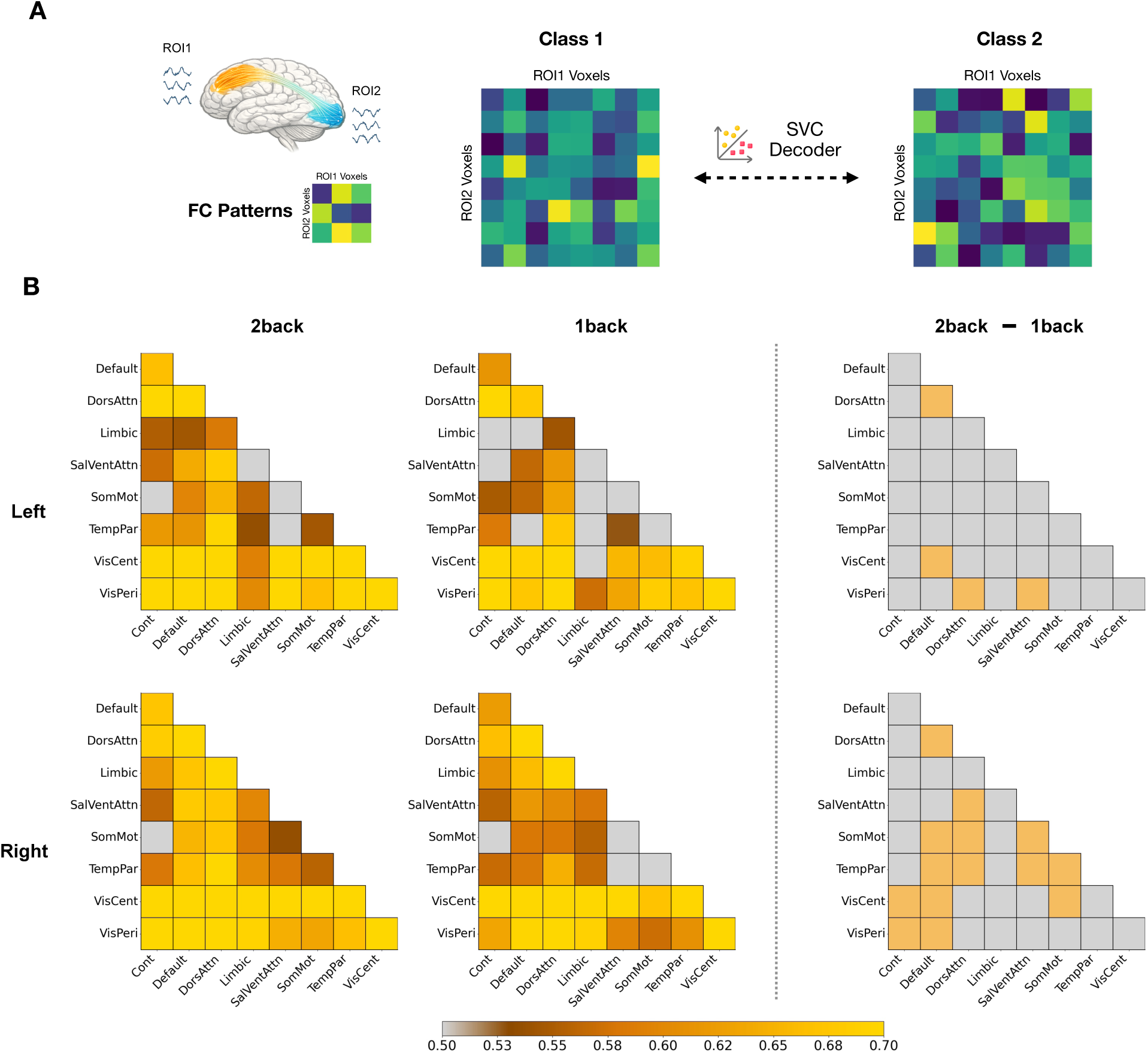
Load Effects Extend Beyond Individual Brain Regions to Inter-Regional Coordination. **A**: Schematic of the FCPA analysis. In this analysis, we compute pairwise correlations between voxel time courses across two ROIs or networks, and use the resulting voxel-wise connectivity patterns to decode visual cognitive information. **B**: FCPA decoding accuracy for object category under different load conditions across pairs of networks in the left and right hemispheres from experiment 1 data. Decoding accuracy for each network pair was tested against chance level (0.5) using one-sample t-tests, followed by FDR correction across all network pairs within the same hemisphere. Contrast maps were generated using one-sided t-tests, followed by FDR correction across the same set of network pairs. Network pairs that did not survive correction are shown in light gray.

With FCPA, we found that, in Experiment 1, the connectivity patterns between various pairs of functional networks indeed enabled significant readout of object, attention, and load (Supplementary Figure 7). An additional analysis confirmed that FCPA decoding is distinct from the underlying univariate activation magnitude or multivariate within-ROI voxel patterns, as regressing out these signals from the functional connectivity patterns did not eliminate the decoding (Supplementary Figure 8). This provides evidence that functional connectivity patterns carry representational information beyond local brain patterns, offering an additional substrate for the widespread representation of information across the brain.

Having demonstrated the validity of FCPA, we next used it to examine if load also modulates the inter-regional representation of object information, paralleling its effects at the level of individual ROIs. Under low load, object categories could be reliably decoded from the connectivity patterns of most network pairs, (Figure 3B; 26/36 pairs left, 32/36 pairs right; p’s < 0.05, FDR-corrected). As load increased, object information became decodable from a broader set of region pairs (32/36 pairs left, 35/36 pairs right; 23.1% and 9.4 % increases, respectively). This more widespread inter-regional object decoding mirrors the MVPA results, where higher load was likewise associated with a broader spatial distribution of object representations. Direct contrasts between the two load conditions revealed significantly enhanced decoding in multiple network pairs (4/36 pairs left, 14/36 pairs right; p’s < 0.05, FDR-corrected). A more stringent analysis focusing on a selected set of ROIs in the control, default, dorsal attention, and visual networks, while regressing out univariate activity and within-region activity patterns, further confirmed the enhanced visual representations under higher cognitive load (Supplementary Figure 9; 1-back: 48/630 pairs left, 18/435 pairs right; 2-back: 172/630 pairs left, 124/435 pairs right; p’s < 0.05, FDR-corrected; 258.3% and 588.9% increases, respectively). The enhanced decoding of object category from functional connectivity patterns at the network level was also replicated using data from experiment 2 (Supplementary Figure 10; 1-back: 30/36 pairs left, 34/36 pairs right; 2-back: 34/36 pairs left, 36/36 pairs right; 11.1% and 5.5% increases, respectively; Paired t-test: 23/36 left, 31/36 right; p’s < 0.05, FDR-corrected). Together, these findings suggest that the brain also leverages stronger inter-regional representations of object information to support visual processing under heightened cognitive demands.

Interestingly, similar to load, attention also modulates the strength of object decoding from multivariate connectivity patterns (Supplementary Figure 11; 12/36 network pairs left, 24/36 network pairs right showed higher object decoding under shape than color attention). This parallels the MVPA results, where shifting attention also produced comparable changes in the spatial extent of object representations (Figure 1C). These findings raise the possibility of a robust relationship between the broader spatial distribution of visual representations and the stronger inter-regional representations, as the latter may facilitate the propagation of object information across distributed cortical regions.

### Load Stabilizes Inter-Trial Multivariate Neural Patterns

In addition to pattern decoding strength, another property that has been largely overlooked in previous fMRI literature on cognitive load is the variability of neural patterns. Previous electrophysiological studies have shown that reduced trial-to-trial neural variability is associated with more efficient response preparation (Churchland et al., 2006, 2010), with similar stabilizing effects also observed in univariate BOLD responses in a limited number of fMRI studies (He, 2013; Ito et al., 2020).

However, none of these studies have examined how variance changes at the level of multivariate brain activity patterns, which serve as an important substrate for representing information. Here, we extended the concept of variance reduction to multivariate neural representations and hypothesized that a similar reduction in variability could also occur in vertex patterns, as heightened cognitive load places greater demands on the brain to process information efficiently. To test this hypothesis, we specifically examined whether increasing cognitive load stabilizes distributed object representations across repeated n-back trial blocks. For each condition, we quantified pattern variability as the mean Euclidean distance between each trial-block pattern and the average pattern across all trial blocks.

Using this approach, we found that higher load was indeed associated with reduced inter-trial variability in object category patterns, indicating more stable object representations under greater cognitive demand. This reduction in variance was significant in almost all brain regions in Experiment 2 (Figure 4A; 96 left, 92 right, p’s < 0.05, FDR-corrected). This global reduction of object pattern variance was similarly observed in Experiment 1, and is present regardless of whether attention is paid to color (Figure 4B; 89 left, 93 right, p’s < 0.05, FDR-corrected) or shape (Figure 4C; 69 left, 76 right, p’s < 0.05, FDR-corrected). Critically, this pattern differs from the load-related expansion of object decoding, which emerged primarily when attention was directed to the object itself. Thus, we conclude that load-related pattern stabilization appears to be a more pervasive, attention-agnostic mechanism that operates commonly on neural representations across the cortex. A similar variance reduction effect was also observed for neural patterns representing attentional states, with higher load significantly reducing pattern variability across widespread cortical regions (Supplementary Figure 3B; 88 left, 92 right; p’s < 0.05, FDR-corrected). This generalizable effect points to a broader influence of cognitive load on visual cognition and supports its overarching role in guiding task-related processing.

**Figure 4.**
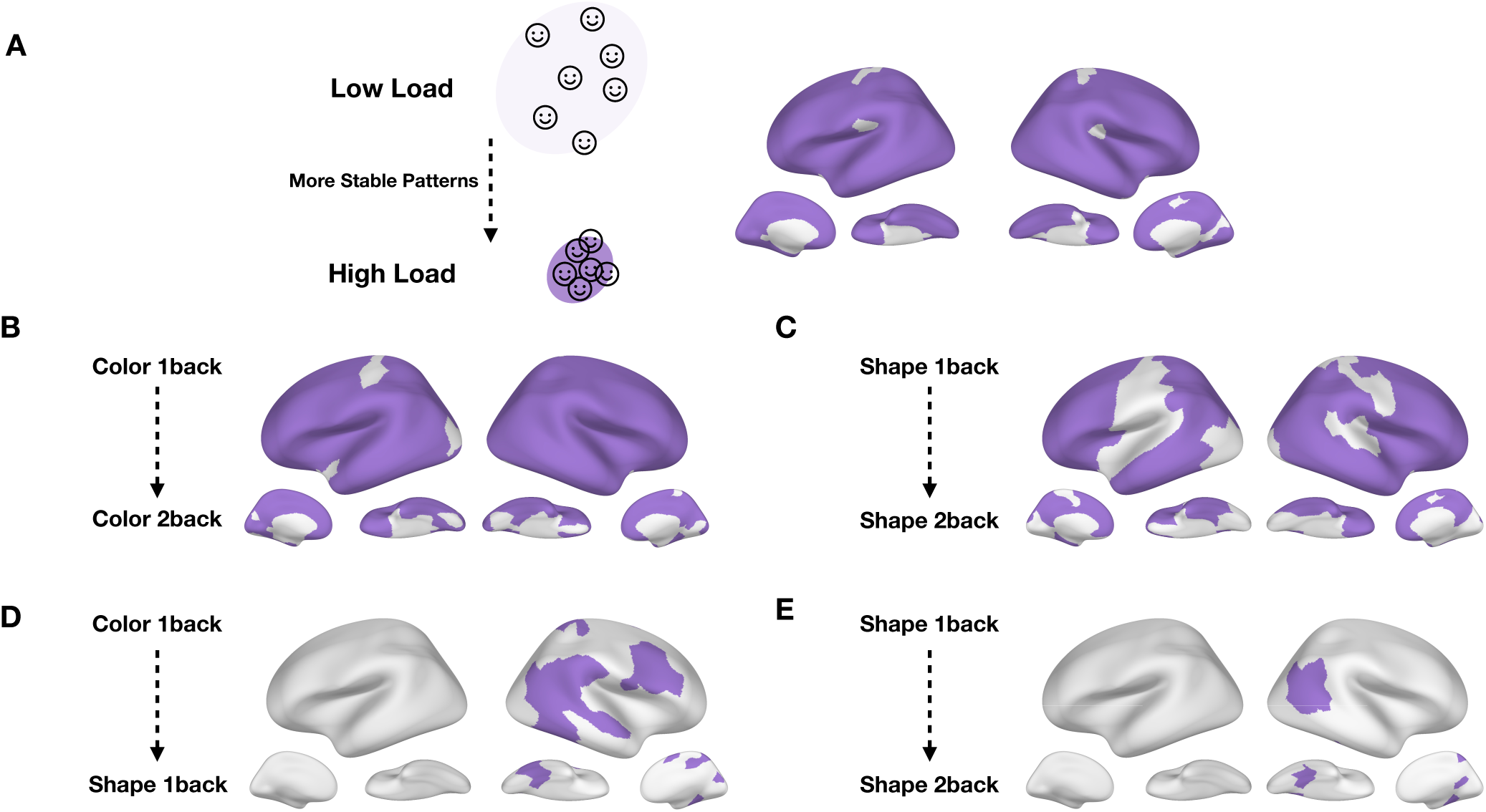
The Variance Reduction Effect on Object Representations. **A**: Cortical regions showing more stable (less variable) object representations under high compared to low load in Experiment 2. Regions with significant reductions in pattern variability (p’s < 0.05, FDR-corrected across all ROIs within each hemisphere) are shown in purple. **B**: Cortical regions showing more stable object representations from color 1back to color 2back conditions in Experiment1 (p’s < 0.05, FDR-corrected). **C**: Cortical regions showing more stable object representations from shape 1back to shape 2back conditions in Experiment1 (p’s < 0.05, FDR-corrected). **D**: Cortical regions showing more stable object representations from color 1back to shape 1back conditions in Experiment1 (p’s < 0.05, FDR-corrected). **E**: Cortical regions showing more stable object representations from color 2back to shape 2back conditions in Experiment1 (p’s < 0.05, FDR-corrected).

One further question is whether this variance reduction is unique to load or reflects a more general strategy used by other task variables, such as attention. In other words, while attention can also strongly modulate representational saliency (Figure 1C), does it do so by similarly reducing pattern variance? When examining changes in inter-trial object pattern variance across attention conditions, in contrast to the effects of increasing load, shifting attention from color to shape did not substantially reduce inter-trial pattern variance. If anything, fewer regions exhibited more stabilized neural patterns when directing attention from color to shape under high load (Figure 4D, E; 1back: 0 left, 24 right; 2back: 0 left, 8 right; p’s < 0.05, FDR-corrected). Together, these findings indicate that the mechanism through which load enhances object representations is not globally shared with other variables, uncovering a unique process by which cognitive load regulates visual processing.

### Load Maintains an Independent Neural Representation

As a key task state that dynamically modulates distributed representations of other task variables, how is cognitive load itself encoded? There are two possible accounts. One possibility is that load is primarily encoded through its influence on other representations, such as the reduction in pattern variability observed in object representations. Alternatively, load may also be encoded as a distinct cognitive state, in addition to its impact on other task variables. To distinguish between these possibilities, we performed cross-object load decoding analyses. As illustrated in Figure 5A, if load is encoded merely through its modulation of object representations, then the neural patterns distinguishing different levels of load should depend on object identity. Consequently, a load decoder trained on some objects should not generalize to other objects. By contrast, if load is also encoded as an independent state, load-induced pattern differences should be largely invariant to object identity, allowing load decoding to generalize across objects with minimal loss in accuracy. Cross-object decoding therefore allows us to distinguish whether load is represented as a separate cognitive state in addition to its modulation of other representations.

**Figure 5.**
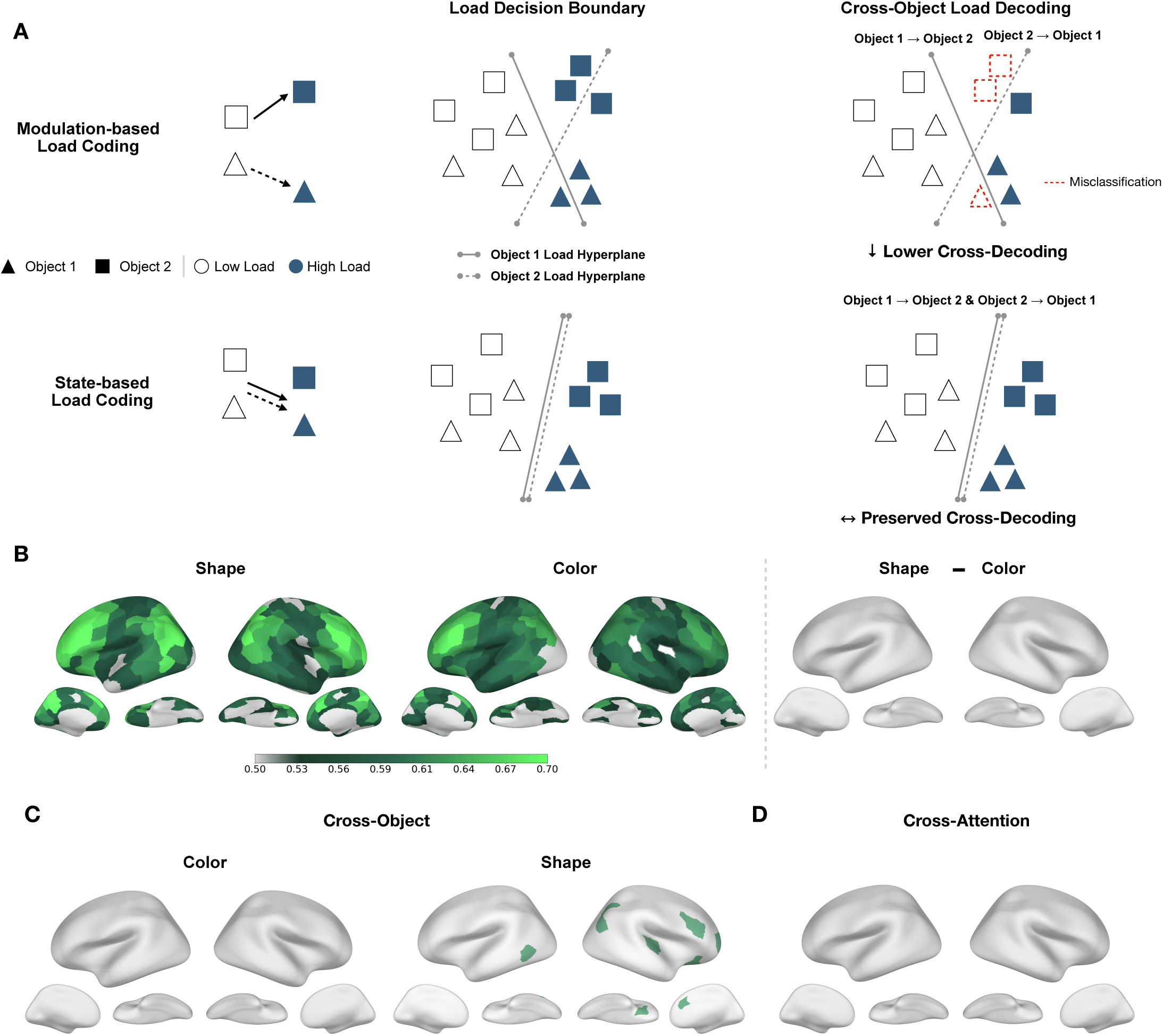
Load Is Globally and Independently Represented Across the Cortex. **A:** Illustration of two potential ways cognitive load could be represented in the brain. Load could be encoded through its modulation of other variables, such as its impact on object representations. Under this account, load-induced pattern changes would be object-specific, resulting in reduced decoding accuracy (i.e., more misclassification) when generalizing across objects. Alternatively, load could be encoded as an independent cognitive state, such that load-induced pattern changes will remain consistent across objects. Under this account, cross-object decoding would show minimal or no drop in accuracy. **B**: Whole-brain decoding maps for load under different attention conditions. Contrast map for load decoding between two attention conditions is shown on the right. Decoding accuracy at each region was tested against chance level (0.5) using one-sample *t*-tests, followed by FDR correction across all ROIs within each hemisphere. Contrast maps were generated using one-sided t-tests followed by the same FDR correction procedure. Regions that did not survive correction are shown in light gray. **C**: Binary decoding drop maps for load across object categories. ROIs shown in green exhibit significantly reduced decoding performance under cross-object decoding (p < 0.05, FDR-corrected), indicating object-dependent load representations. Regions not surviving correction are shown in light gray. **D**: Binary decoding drop maps for load across attention conditions.

Before performing the cross-decoding analysis, we first examined where load could be reliably decoded, specifically whether its representation is localized to the canonical frontoparietal control regions or extends well beyond these regions, as suggested by recent studies on load and arousal representations (Adam et al., 2025; Raut et al., 2025). Reinforcing its global modulatory role, we found that load information was well decodable with MVPA in nearly all cortical regions, with the exception of a few areas in the ventral occipital and medial frontal cortex (Figure 5B, Supplementary Figure 2C; Experiment 1: Color, 89 left, 84 right; Shape, 89 left, 89 right; Experiment 2: 98 left, 100 right). This widespread pattern coding of cognitive load supports the notion that load signals are globally distributed, potentially serving as a regulatory signal that guides the operations of individual brain regions, rather than being confined to higher-level cognitive regions.

When decoding cognitive load across object categories, we found that across most of the cortex, neural decoding of cognitive load remained largely unaffected, suggesting that load is encoded more as a separate cognitive construct in addition to its modulatory effects on other representations (Figure 5C). Here we observed only a few scattered regions that showed drops in decoding accuracy in the shape task (1 left, 7 right). These regions, however, did not replicate when we included eight categories in Experiment 2, where no regions exhibited a cross-object load decoding drop (Supplementary Figure 12). Overall, these results are in support of the coding of load as an independent cognitive state. This stands in contrast with the other task variable, attention, which displays more widespread cross-object decoding drops, especially under higher load (Supplementary Figure 13, 1back: 5 left, 0 right; 2back: 41 left, 22 right).

We also demonstrated that load coding is relatively stable across the other task variable, the attentional states. When directly contrasting the two attention conditions, no regions showed significant changes in load decoding accuracy (Figure 5B). Consistent with this invariance in decoding strength, when decoding load across attention conditions, no regions exhibited significant cross-decoding drops (Figure 5D), further consolidating the notion that the neural coding of load is minimally influenced by the attention deployed. Lastly, examining the inter-trial pattern variability of load representations (Supplementary Figure 14), no left hemisphere regions showed more stable neural patterns when attention shifted from color to shape, while 37 of 100 right hemisphere regions exhibited such a reduction in variance. This effect was considerably more restricted than the widespread variance reductions induced by higher load on attention representations (Supplementary Figure 3B; 100% and 59.8% decreases, respectively, for left and right hemispheres), implying that attention exerts only a limited influence on the stability of load representations.

Overall, these results established a distinct representational scheme for cognitive load: as a task signal that broadly modulates other visual cognitive representations, cognitive load is encoded throughout the entire cortex, while its own neural coding remains largely unaffected by other visual and task information. This coding scheme suggests that load is widely represented as an independent cognitive state in addition to its ability to directly modulate neural patterns, reinforcing its role as an overarching control signal during visual cognitive processing.

## Discussion

Using n-back fMRI data, our work demonstrated that high cognitive load is accompanied by enhanced neural representations of visual cognitive information across the brain, potentially supporting successful task performance despite increased cognitive demands. Specifically, we identified four important properties of cognitive load during visual processing. First, higher load was associated with stronger and more distributed representations of visual information. Second, the pervasive effect of load extended beyond individual brain regions to shape and coordinate visual representations across and between multiple brain networks, as reflected in the increased representational alignment across cortical networks and more salient decoding from inter-regional co-activation patterns under high load. Third, under heightened cognitive demands, the brain represented visual cognitive information with more stabilized, less variable neural patterns. Finally, as a modulatory signal, cognitive load itself was encoded in a widely distributed fashion across the brain as a separate cognitive state, with its neural representation remaining largely independent of other visual and task variables.

### Enhanced and Widespread Visual Representation Under High Load

Across the two experiments, we found that higher load gives rise to stronger and more widespread decoding across the brain, beyond what has been shown previously. This enhancement of visual representations under heightened cognitive demands was observed in the lPFC, PPC, and OTC (Shao et al., 2025; Xu & Chun, 2025). Importantly, our findings extend previous work by demonstrating that such enhancement is not restricted to these canonical regions, but instead encompasses the majority of the cortex.

This enhancing effect of load diverges from the traditional account that higher cognitive load primarily impairs visual cognition (de Fockert et al., 2001; Kelley & Lavie, 2011; Lavie & De Fockert, 2005; Todd et al., 2005). We believe that these findings are not necessarily contradictory. While higher cognitive load could result in performance decline (Figure 1B, Supplementary Figure 1C, 2B), the brain may attempt to counteract this challenge by recruiting additional neural resources and reorganizing representations to maximally preserve salient traces of task-relevant information. In this sense, the stronger and more widespread representation of visual cognitive information may reflect an adaptive response to increasing cognitive demands. However, because the neural resources available to support task performance are ultimately finite, such compensatory increases in neural representation may become insufficient once cognitive demands exceed the available capacity, resulting in the behavioral impairments typically associated with high load. In addition, our decoding analyses focused on higher-level object category and attentional state information rather than item-specific information, the latter of which has been reported to deteriorate under higher load (Chopurian et al., 2026; Emrich et al., 2013; Sprague et al., 2014). This discrepancy may imply that under greater cognitive demand, increased neural effort preferentially preserves task-relevant information that is stable and shared across trials, while lower-level, item-specific features are represented with reduced precision. Thus, rather than uniformly enhancing or degrading visual representations, cognitive load may reshape how limited neural resources are allocated across different levels of visual information.

Our finding of more distributed visual cognitive representations aligns with and extends previous ROI-based findings that visual information is represented in regions beyond the ventral visual pathway (Konen & Kastner, 2008; Vaziri-Pashkam & Xu, 2017). Moreover, it is consistent with findings from rodent studies showing that increasing computational demands similarly drive more distributed encoding of task-relevant variables (Rosen & Freedman, 2025). This cross-species convergence suggests a fundamental adaptive strategy, whereby the brain coordinates additional neural resources beyond canonical visual regions to support task-relevant information under increasingly complex and demanding environments.

To date, distributed visual cognitive representations remain relatively underexplored in tightly controlled laboratory fMRI paradigms, where much of the literature has instead focused on canonical regions associated with specific visual functions. Our finding that cognitive load can induce more widespread visual representations suggests that the neural substrates supporting visual cognition are more dynamic and adaptive than traditionally characterized. In addition, our results may provide a bridge between the localized representation elicited by controlled simple lab tasks and the widespread representations extensively observed under naturalistic settings (Çukur et al., 2013; Huth et al., 2016). We hope this line of work can motivate further investigation into the dynamic processes through which visual information is represented across the brain.

### High Load Aligns Cortical Representational Structures

The impact of cognitive load on visual representations extends beyond individual brain regions to the organization of information across the broader cortex. Our results showed that higher load promotes greater convergence in how visual information is represented across different brain networks.

This large-scale alignment implies that, as cognitive demand increases, the brain may shift toward a more coherent mode of processing task-relevant information, rather than relying on a more segregated division of labor across specialized regions. We speculated that this convergence reflects another adaptive response to increased task demands: by bringing representations across networks into greater alignment, the brain may facilitate coordinated processing of task-relevant information and provide additional safeguards for its preservation and subsequent use under demanding conditions. Such an account is consistent with recent findings from non-human primates (Liu et al., 2026) and computational models (Doimo et al., 2022).

Notably, the increase in representational alignment cannot be explained solely by the greater number of brain regions exhibiting significant visual information decoding under high load. Instead, the increased alignment remained evident even after excluding regions that only showed significant decoding under high load (Supplementary Figure 6D-H). This indicates that load does more than simply expand visual representations into additional cortical regions; it also reorganizes representational structures that are already present across the cortex.

This global representational change in brain representation also aligns with previous work that probed cognitive load at the broader network level. Graph network analysis of both fMRI and EEG data (J. R. Cohen & D’Esposito, 2016; Kitzbichler et al., 2011; Shine et al., 2016) suggested that when the brain is exposed to higher cognitive demands, multiple brain regions showed heightened synchronization in their temporal activity profiles. However, these functional connectivity measures primarily captured statistical associations between activity fluctuations across brain regions, leaving the functional significance of such coordinated activity largely unexplained. Moreover, they did not directly characterize the multivariate activity patterns that are critical for understanding how information is represented in the brain.

Our findings help bridge this gap by demonstrating that higher cognitive load is accompanied by a global reorganization of representational structure, such that distributed brain regions encode visual information in increasingly similar formats. This shift toward representational alignment suggests that coordinated brain activity may facilitate the transmission and integration of shared information across distributed cortical regions, beyond localized processing. Future studies can further characterize how functional connectivity and representational alignment are mechanistically linked.

While higher load promotes representational convergence, it does not completely eliminate the differences in representational structure preserved across different brain networks, as evident in Figure 3C. Canonical visual regions, for instance, may still represent object information in a different format from frontoparietal control regions, even as their overall representational structures become more aligned under higher load. We therefore propose that representational convergence and specialization are not mutually exclusive forms of cortical organization, but can coexist to varying degrees, with their relative balance dynamically shifting according to task demands. Under higher cognitive demand, though, this balance may shift toward greater convergence as an adaptive mechanism to optimize cognitive processing and behavioral performance.

### Load Modulates Interregional Representational Strength

Using FCPA, we showed that higher cognitive load not only enhances information readout within individual brain regions but also improves the decodability of information from a novel representational substrate, namely, the functional connectivity patterns between pairs of brain regions.

Importantly, FCPA offers a novel approach to characterizing information encoded in functional connectivity patterns. Unlike existing multivariate connectivity-based approaches that often involve dimensionality reduction or infer representational information indirectly from transformed connectivity spaces, FCPA directly leverages raw voxel-to-voxel connectivity strength patterns for decoding. This approach minimizes data transformations while providing a framework closely analogous to conventional MVPA, extending multivariate decoding from local activity patterns to interregional connectivity patterns. Critically, by controlling for both univariate activity and multivariate within-ROI representations (Supplementary Figure 8, 9), we further demonstrated that functional connectivity patterns carry additional representational information that cannot be entirely explained by these more conventional forms of neural signals. Together, these findings highlight not only the increased contribution of interregional representations under high cognitive load, but also FCPA’s potential as a complementary approach for uncovering information encoded in distributed neural interactions.

This enhanced connectivity-based representation, together with the increased alignment of representational structures across cortical regions, provides converging evidence that the effect of load on visual processing is not localized to individual regions, but instead manifests as a global property across the brain. We speculate that these two effects may be inherently linked, whereby more salient interregional representations under high load may facilitate the propagation of visual information across cortical regions, ultimately contributing to greater representational convergence across the brain.

We hope these findings draw greater attention to the representational power embedded in multivariate functional connectivity patterns, which has received little consideration compared to the univariate magnitude of connectivity strength. Future research could build upon this framework to systematically characterize the correspondence and dissociation between regional and inter-regional representations across different cognitive domains, as well as the computational significance of representing information through these distinct forms of neural patterns.

### Load Stabilizes Visual Representation Across The Brain

In addition to enhancing visual representation, we demonstrated for the first time that load also largely reduces trial-to-trial variability in neural patterns. Trial-to-trial variance in neural signals has been primarily studied using electrophysiological recordings in humans and non-human primates, where the term “neural quenching” describes the sharp reduction in signal variability following perceptual stimulus onset (Arazi, Gonen-Yaacovi, et al., 2017; Churchland et al., 2010; M. R. Cohen & Maunsell, 2009; Daniel et al., 2019; Terlau et al., 2026). Such reductions in signal-amplitude variability have been proposed to reflect greater certainty in information representation (Goris et al., 2024), and are associated with individual behavioral performance (Arazi, Censor, et al., 2017; He, 2013; Samanez-Larkin et al., 2010).

Despite its potential importance, variance reduction in univariate neural activity has received relatively little attention in the fMRI literature (He, 2013; Ito et al., 2020). Even less is known about whether such a stabilizing effect extends to multivariate activity patterns, which carry rich information about neural representations. Here, we extended this finding to the level of population patterns using fMRI data and found that higher cognitive load consistently reduced pattern variance, regardless of the attended features. This attention-agnostic effect partially dissociates variance reduction from enhanced decoding, as the latter was observed only when attention was directed toward the objects. Such a distinction suggests that variance reduction may reflect a more general mechanism the brain uses to cope with increased cognitive demands. In other words, to support efficient and accurate processing under heightened load, the brain may prioritize maintaining cleaner and more stable neural representations that capture the essential information being processed at a given moment.

### Load as a Standalone Modulatory Signal

While load modulated visual cognitive representations, the neural representation of load itself remained relatively insensitive to changes in other task variables, as indicated by both within- and cross-decoding results. In particular, the absence of a cross-object decoding drop for load further suggests that load is not encoded merely as a byproduct of its variance-reduction effect on object representations. Instead, load appears to be encoded as an independent task state in addition to its modulatory effect on other variables, and it is minimally susceptible to changes in other task variables.

This independence implies that load serves as an overarching task factor in visual cognition that exerts broad influence over other representations while remaining largely insulated from them. Such representational independence could be critical for load to continuously regulate ongoing processing across different task contexts. This account aligns with interference-resolving mechanisms previously demonstrated in both neural data and computational models, where orthogonalized representations help different task variables to co-exist flexibly (Bernardi et al., 2020; Flesch et al., 2022; Weber et al., 2023).

The modulatory role of load is further underscored by the observation of widespread load decoding across cortical regions, as well as by FCPA decoding between various networks. Notably, our findings substantially extend previous work by demonstrating that representations of cognitive load are distributed far beyond the traditional frontoparietal or multiple-demand networks (Duncan, 2010; Marek & Dosenbach, 2018), suggesting that load may permeate more broadly across the brain to support task monitoring and adaptive cognition.

Overall, across two experiments, we provide converging evidence for how the brain dynamically adapts to increasing cognitive demands. Rather than simply deteriorating under higher cognitive load, the brain actively responds to these demands by recruiting more widespread neural resources, coordinating and promoting representations across brain regions, stabilizing multivariate activity patterns, and maintaining independent representations of cognitive load, to potentially support more efficient information processing. These findings highlight the remarkable flexibility of neural representations in adapting to changing task demands (Bracci et al., 2017; Kay et al., 2023; Nau et al., 2024), and underscore the importance of understanding brain function and cognition from a more dynamic perspective that extends beyond localized neural processing (Rosen & Freedman, 2025).

## Methods

### Data Collection and Participants

Experiment 1 conducted new analyses of data previously collected and published by Xu & Chun (2025). Twelve healthy, right-handed adults (8 females, 4 males; age range: 18–35 years) with normal or corrected-to-normal vision participated in the study.

The fMRI data for Experiment 2 were newly collected. Twelve healthy, right-handed adults (8 females, 4 males; age range: 18–35 years) with normal or corrected-to-normal vision participated. One participant (female) took part in both experiments. All participants provided written informed consent prior to participation and received monetary compensation. Both studies were approved by the Committee on the Use of Human Subjects at Yale University.

### Experiment 1 Design

Experiment 1 is a visual n-back task that allows manipulation of the three variables of interest: object category, attention, and task load. In each n-back trial (Figure 1A, Supplementary Figure 1A, B), ten images were presented sequentially (200 ms each), each followed by a 600 ms display of a red fixation dot at the center of the screen. Participants were instructed to maintain fixation on the central red dot throughout each trial. The ten images presented in a given trial were different exemplars of the same object category. The object categories included cars, cats, faces, and houses, chosen to represent real-world objects commonly encountered in everyday life. These stimuli have been validated and used in prior studies (Vaziri-Pashkam et al., 2019; Vaziri-Pashkam & Xu, 2017, 2019; Xu, 2023; Xu & Vaziri-Pashkam, 2019). For each category, ten unique exemplars were created, with variations in viewing angle, identity, pose (for cats), and facial expression (for faces). Some exemplars are shown in Supplementary Figure 1B. As in Experiment 2 of Vaziri-Pashkam & Xu (2017), in addition to the grayscale image itself, each stimulus was superimposed with 30 semi-transparent colored dots that followed the same spatial envelope of the original object. The 30 dots in each image had the same color, randomly chosen from the ten different colors used throughout the experiment.

As noted above, each trial block consisted of 10 images interleaved with red fixation dots and lasted 8 seconds. Between consecutive trial blocks, 8-second fixation blocks were presented, during which participants were instructed to simply maintain fixation without performing any task. Among the ten images presented in each trial, two were randomly selected to be the n-back repeats.

Using an n-back paradigm coupled with color-dotted object images allows us to manipulate both task load and attended features. Specifically, the experiment included two load conditions (1-back and 2-back) and two attention conditions (attending to color vs. shape), resulting in four (2-by-2) combinations of task conditions (Figure 1A, Supplementary Figure 1A). Sample correct responses for each of the four task conditions are shown in Figure 1A. In the fMRI session, each participant completed sixteen runs, where each of the four task conditions occupied four runs. To avoid ordering effects and ensure that certain task conditions did not consistently appear earlier than others, we balanced the task order as follows: for odd-numbered participants, the task order for each run was fixed as 1, 2, 3, 4, 3, 1, 4, 2, 2, 4, 1, 3, 4, 3, 2, 1. For even-numbered participants, the order was 2, 4, 1, 3, 4, 3, 2, 1, 1, 2, 3, 4, 3, 1, 4, 2. The numbers 1–4 correspond to the four task conditions: color 1-back (1), color 2-back (2), shape 1-back (3), and shape 2-back (4), respectively. Within each run, each object category appeared four times, tallying up the sixteen trial blocks. The presentation order of the category was determined by a balanced Latin square.

To minimize task-switching effects across runs, we included an additional practice trial block at the beginning of each run (not known by the participant). Data from this trial block were excluded from analysis. As a result, each run contained 17 trial blocks, each lasting 8 seconds. Combined with sixteen interleaved 8-second fixation blocks, a 2.4-second initial fixation period, and an 8-second final fixation period, each run lasted a total of 4 minutes and 34.4 seconds.

### Experiment 2 Design

To better characterize the fine-grained object representational structures in different brain regions across loads, we conducted Experiment 2 with an increased number of object categories (Supplementary Figure 2A). The eight object categories used here were cars, cats, houses, faces, bodies, chairs, elephants, and scissors. The stimuli were introduced in Vaziri-Pashkam & Xu (2017) and adapted here. As in Experiment 1, cognitive load was manipulated using separate runs of 1-back and 2-back tasks. In contrast to Experiment 1, there was no attention manipulation; thus, the grayscale object images were presented directly without any color dot overlay.

Sixteen runs of data were collected from each participant, with each of the two task conditions occurring in eight runs. A pseudo-random run order was pre-set for each participant as follows: for odd-numbered participants, the run order was 1, 1, 2, 2, 1, 1, 2, 2, 2, 2, 1, 1, 2, 2, 1, 1; for even-numbered participants, the run order was 2, 2, 1, 1, 2, 2, 1, 1, 1, 1, 2, 2, 1, 1, 2, 2. Here, 1 denotes the 1-back task, and 2 denotes the 2-back task. Within each run, each object category appeared twice, resulting in sixteen trials per run. All other experimental details followed those outlined above for Experiment 1.

### MRI Acquisition

#### Experiment 1

The following MRI acquisition details are reproduced and summarized from Xu & Chun (2025). Participants underwent two scanning sessions, each lasting approximately 1.5 hours, which included both a localizer task and the main experiment. Data were collected using a Siemens Prisma 3T scanner at the Yale Brain Imaging Center equipped with a 32-channel receiver head coil. High-resolution T1-weighted structural images (0.8 × 0.8 × 0.8 mm) were acquired using an MP-RAGE sequence (Mugler III & Brookeman, 1990). Functional images were collected using a T2*-weighted echo-planar imaging (EPI) sequence that incorporated multiband RF pulses and simultaneous multi-slice (SMS) acquisition. Whole-brain coverage was achieved with 72 axial slices at 2 mm isotropic resolution (TR = 800 ms, TE = 37 ms, flip angle = 52°, SMS factor = 8).

#### Experiment 2

The data in this experiment were collected at BrainWorks at the Center for Neurocognition and Behavior in the Wu Tsai Institute, Yale University (RRID: SCR_024556), using a Siemens Prisma 3T scanner equipped with a 32-channel receiver head coil. All other parameters are the same as in Experiment 1.

### MRI Preprocessing

The data were primarily preprocessed using FreeSurfer (Fischl, 2012) and FsFast (Dale et al., 1999). The preprocessing pipeline mainly included 3D motion correction as well as linear and quadratic trend removal. Functional data were then co-registered to the high-resolution anatomical images acquired with an MP-RAGE sequence and projected onto each participant’s native inflated cortical surface. All the following fMRI analysis in this study are performed based on the surface-based data, given its higher sensitivity (Oosterhof et al., 2011) and lower susceptibility to noise (Brodoehl et al., 2020) over volumetric data.

### fMRI Analysis

#### ROI Definition

The whole brain was parcellated into 200 regions of interest (ROIs) using the Schaefer 200-parcel atlas (Schaefer et al., 2018), with 100 ROIs in each hemisphere. The atlas was projected from fsaverage space onto each participant’s native surface using the FreeSurfer command *mri_surf2surf*. All subsequent multivariate pattern analyses (MVPA) were conducted separately within each ROI, and the corresponding results are mostly visualized by various colored brain maps in the following sections. For analyses focusing on selected ROIs, we extracted data using the corresponding parcels from the Schaefer atlas in the same way as we did the whole-brain analysis. For network-level analyses, we used the 17-network version of the Schaefer atlas, in which each ROI is assigned to one of 17 canonical functional networks. Networks with similar labels (e.g., Visual A and Visual B) were further combined into the broader networks (e.g., Visual) for more concise interpretation.

#### MVPA Within-Condition Decoding

Multivoxel patterns from each ROI come from the univariate response magnitudes of individual vertices within that ROI. We used a general linear model (GLM) to estimate beta coefficients that reflect the response magnitude. The GLM is applied independently to each run for every participant. Within each run, each 8-s trial block was modeled as a separate event, yielding one beta estimate per block. The resulting activity patterns for each trial block were then z-normalized to preserve relative magnitude differences across vertices while removing trial- or run-specific amplitude differences. Taken together, for Experiment 1, this yielded normalized vertex activity patterns within each ROI for every trial block of the 16 experimental conditions (2 attention × 2 load × 4 object categories). For Experiment 2, the same procedure again yielded normalized multivariate activity patterns within each ROI for every trial block of the 16 experimental conditions (2 load × 8 object categories).

With these patterns in hand, we proceeded with the multi-voxel pattern analysis (MVPA) for Object, Load, and Attention in Experiment 1, and for Object and Load in Experiment 2. We implemented MVPA here using customized Python scripts and used the SVC function (with a linear kernel) from the scikit-learn package (Pedregosa et al., 2011) to perform the classification. For both Experiment 1 and 2, we performed binary classification between each pair of the 16 conditions (Experiment 1: 4 object categories x 2 attention x 2 load; Experiment 2: 8 object categories x 2 load) and utilized a leave-one-run-out cross-validation procedure to improve the reliability of our results. The procedure works as follows: for each of the 16 conditions, we obtain 16 pattern vectors corresponding to the 16 trials (4 runs × 4 trials per run) associated with that condition. In each iteration of the 4-fold cross-validation, one run is held out as the test set, while the remaining three runs serve as the training set. Classification is performed on the training data, and the procedure is repeated four times so that each run serves as the test set once. The final classification accuracy is computed by averaging the results across all four iterations. To ensure that differences in vertex count did not confound decoding accuracy, we controlled the number of vertices used for decoding across ROIs: In each cross-validation iteration, the top 500 vertices were selected based on paired t-statistics contrasting the two decoded conditions, computed exclusively from the training data.

From the 16-by-16 classification accuracy matrix, we can derive the decoding strength of a given variable while accounting for the other variables. For instance, in Experiment 1, to assess object decoding under the shape 2-back condition, we averaged the pairwise decoding accuracies between all pairs of object categories within the shape 2-back task. That is, the average MVPA accuracy across all pairwise decoding of the following four conditions: “shape 2-back, cars”, “shape 2-back, cats”, “shape 2-back, houses”, and “shape 2-back, faces”. Similarly, we derived the object decoding under the shape 1-back condition by averaging the pairwise decoding accuracies across the four object categories within the shape 1-back task. To subsequently assess load-dependent enhancement of object representations when attending to shape, we performed paired t-tests comparing the resulting object decoding maps from the shape 1-back and shape 2-back conditions, followed by FDR correction for multiple comparisons.

All analyses were conducted independently within each ROI defined in the previous section (see <u>ROI Definition</u>) for each participant. Group-level decoding performance was evaluated using one-sided t-tests against chance level (0.5), followed by false discovery rate (FDR) correction for multiple comparisons (Benjamini & Hochberg, 1995).

#### MVPA Cross-Condition Decoding

We conducted cross-condition MVPA decoding (hereafter abbreviated as cross-decoding) to examine the interaction and independence of neural pattern representations across variables. This analysis followed the same cross-validation procedure as the within-condition MVPA, except that the training and testing sets were drawn from different conditions rather than the same condition. Specifically, we decoded one variable across levels of another variable while holding the third variable constant. For example, in Experiment 1, to test whether load representations generalized across object categories when attention was directed to color, we trained SVC classifiers to distinguish between low and high load using neural patterns from trials of one object category under the “attending to color” condition, and then tested the classifiers on the corresponding patterns from another object category under the same attention condition. We then repeated the analysis in the opposite direction (i.e., training on the second object category and testing on the first) and averaged decoding accuracies across the two directions to obtain the final cross-decoding performance.

Paired t-tests were performed between the cross-condition decoding and the within-condition decoding performance of a variable, followed by FDR correction. A significant decrease in cross-decoding suggests that the representation is not invariant across the other variable, indicating interactive rather than independent encoding.

#### Representational Structure Analysis

In Experiment 2, with the increased number of object categories, we analyzed the fine-grained representational structures encoded by each ROI’s fMRI activity patterns. This analysis used the pairwise decoding accuracies to construct representational distance matrices (RDMs). Unless otherwise specified, all representational structure analyses below were performed using the group-averaged RDMs across participants.

First, we examined the representational structure of all 16 object (8) × load (2) conditions within a subset of ROIs and networks (Figure 2B, Supplementary Figure 5). Specifically, using the 16 × 16 RDM, we first subtracted the chance decoding accuracy (0.5) from each entry so that only above-chance representational distances were retained. We then applied multidimensional scaling (MDS) to project the RDM into a two-dimensional space while preserving their relative dissimilarities.

To further examine how object representational structures changed between loads across the brain, we then characterized the large-scale organization of object representations across the cortex using a second-order RSA, by constructing an RDM capturing the representational dissimilarity between each pair of ROI (or network) × load combinations (Figure 2A). Specifically, the distance between each pair of combinations was calculated as one minus the Pearson correlation between their corresponding object RDMs. The resulting RDM was then projected into a two-dimensional space using MDS to visualize the representational relationships among ROIs (or networks) under different load conditions. The two-dimensional solution was chosen both for ease of visualization and because it provided an optimal fit to the data (Supplementary Figure 6I, J). The above analysis was conducted at both the ROI and network levels. For the latter, the object RDM for each network was derived by averaging the object RDMs of its constituent ROIs.

To quantify changes in network-level object representational structures across load conditions, we used two measures: the mean distance to the centroid in the MDS space and the mean network RDM correlation. The former is calculated from the MDS coordinates by first computing the center coordinate across all networks, then calculating each network’s Euclidean distance to this center and averaging these distances across networks. This serves as a measure of the overall distance of network-level representation, with smaller values indicating that the object representation of different networks is more closely aligned in the shared representational space. The latter was calculated as the mean pairwise correlation between network RDMs within each load condition, with higher values indicating greater convergence of object representations across networks. Notably, this measure was calculated separately for each participant, allowing a paired t-test to assess whether the effect was consistent across participants.

#### Functional Connectivity Pattern Analysis

While individual ROIs use population-level patterns to represent key variables, it is unlikely that each ROI operates in isolation. Information may also be conveyed through interactions between regions—specifically, in the patterns of communication between ROIs. To test this idea, we developed a procedure termed functional connectivity pattern analysis (FCPA). FCPA closely mirrors the standard MVPA pipeline, but instead of decoding information from within-ROI vertex patterns, FCPA decodes information using vertex-wise functional connectivity patterns between pairs of ROIs.

To increase the reliability of the functional connectivity estimates, rather than computing connectivity separately for each trial (which contained only 10 TRs), we concatenated the time courses from every four trials blocks (with inter-trial fixation TRs removed) into a single time course before calculating functional connectivity. Specifically, within each run, the four trials belonging to the same object category were concatenated to form one sample for the FCPA analysis. As a result, to keep the number of samples equal to the one used in the MVPA analysis, we collapsed across task variables that were orthogonal to the variable being decoded. That is, for object decoding, we pooled trials across all task conditions for each object category before classifying between object pairs. For attention and load decoding, we pooled trials across all object categories for training and testing.

FCPA was performed for all vertices between each pair of ROIs. If ROI A contains M vertices and ROI B contains N vertices as input for a standard MVPA, then each ROI contains M X N edges for the FCPA. To control for the different number of vertices in each ROI, we selected the top 500 vertices in every ROIs based on paired t-tests on activation amplitude (i.e., beta values) between the two decoded conditions, as in MVPA. For each trial, we then computed vertex-wise functional connectivity by calculating Pearson correlations between the time courses of all vertex pairs across the two ROIs, yielding a functional connectivity pattern matrix (in the shape of 500 x 500) for that trial. These patterns were subsequently z-normalized to ensure equal mean and variance. Finally, trials were partitioned into four folds, with three folds used for training and the remaining fold reserved for testing, following the same cross-validation procedure as in the standard MVPA. FCPA was conducted independently between each pair of ROIs selected for each participant. Group-level decoding performance was evaluated using one-sided t-tests against chance level of 0.5, followed by FDR correction for multiple comparisons.

A follow-up analysis was conducted to assess whether FCPA decoding was entirely driven by multivariate vertex patterns within each ROI or by univariate activity differences between conditions. To test these possibilities, we repeated the FCPA analysis described above, but selected the bottom 500 vertices from each ROI based on paired *t*-tests of activation amplitudes (i.e., beta values) between the two decoded conditions. These vertices did not exhibit much amplitude differences between conditions and therefore contained minimal univariate discriminative information. In addition, after computing the functional connectivity pattern matrix for each trial, we regressed out the constituent ROIs’ vertex activity patterns from the connectivity pattern. Because every four trial blocks were concatenated to form a single trial in the FCPA analysis, the vertex patterns from all four constituent trial blocks were included as regressors. All remaining FCPA procedures followed those described above.

#### Inter-Trial Pattern Variance Analysis

Previous studies of neural variability in fMRI have primarily focused on univariate activity (He, 2013; Ito et al., 2020). Here, we extended this approach to multivariate vertex patterns to investigate whether variance reduction also occurs at the level of neural pattern representations. Specifically, we first extracted the trial-wise vertex patterns within each ROI and z-normalized each pattern across vertices. Pattern variability was then quantified as the mean Euclidean distance between each trial’s pattern and the averaged pattern. For object category pattern variance, we first calculated the variance for each object category separately within each task condition (4 in Experiment 1, 2 in Experiment 2), and then averaged across object categories to obtain the overall object pattern variance for each condition. For load and attention pattern variance, we instead pooled trials across all object categories before calculating the variance separately for each condition. To obtain a summary measure for each variable, we then averaged across the corresponding conditions. For example, load pattern variance under the shape condition was calculated by averaging the variance of the shape 1-back trials and the shape 2-back trials.

## Acknowledgement

Research reported in this publication was supported by the National Eye Institute of the National Institutes of Health under Award Number R01EY030854 to YX. The content is solely the responsibility of the authors and does not necessarily represent the official views of the funding agency. The fMRI studies conducted were supported by BrainWorks at the Center for Neurocognition and Behavior in the Wu Tsai Institute, Yale University. We thank Maryam Vaziri-Pashkam for sharing the object stimuli used in the study, and Boyang Hu for his assistance in fMRI data collection.

## Code Availability

Custom Python scripts used for the core analyses in this study, including multivariate pattern analysis (MVPA) and functional connectivity pattern analysis (FCPA), are publicly available on GitHub at https://github.com/howieha/Load-And-Neural-Representation.

## Declaration of Generative AI Use

Generative AI tools were used to assist with manuscript preparation and code development. All scientific content and initial manuscript drafts were written by the authors, with AI tools used to improve clarity, phrasing, and readability. Similarly, the core analysis codebase was developed by the authors, while AI-assisted coding tools were used to help debug, refine, and organize existing code. All AI-assisted modifications were reviewed and verified by the authors, who take full responsibility for the final manuscript and analyses.

---

**Supplementary Figure 1.**
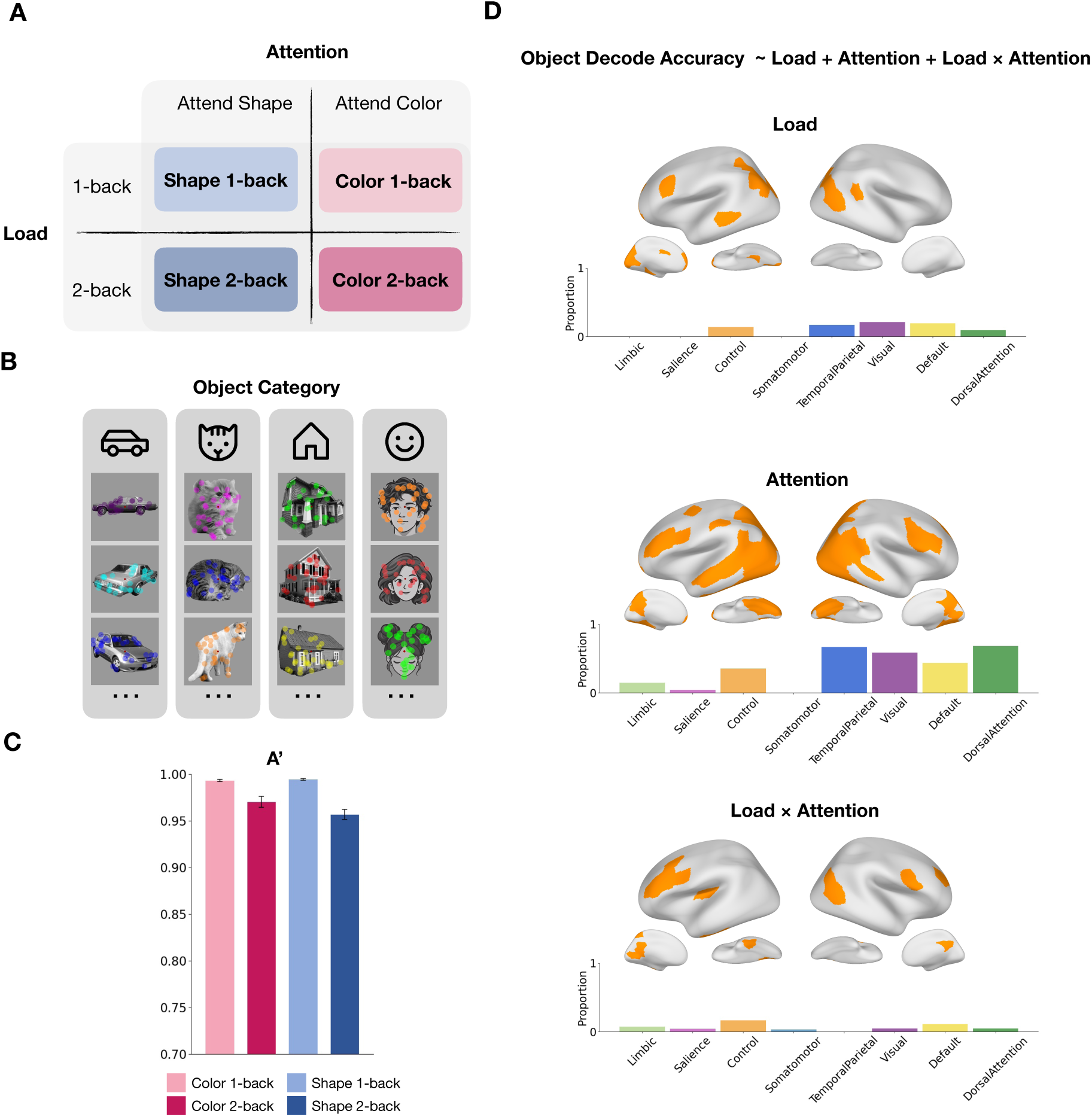
Task Design and Object Category Decoding. **A**: Schematic of a 2×2 factorial experiment design for the n-back task. Each task condition contains a certain attention command (directing to shape versus color) and working memory load (1-back vs 2-back). **B**: The four categories of objects where the trial stimuli come from. Some examples are provided for each object category. The cartoon faces shown here were generated using ChatGPT for illustration; the actual experiment used photographs of real human faces. **C**: A’ behavioral performance under each of the four task conditions (n=12). **D**: ANOVA results for object category decoding. Cortical regions showing significant main effects of attention, load, and their interaction in a linear model predicting decoding accuracy. Only regions surviving FDR correction (p < 0.05) are shown; nonsignificant regions are displayed in light gray. Bar plots below each brain summarize the proportion of regions within each canonical network exhibiting a significant main effect or interaction.

**Supplementary Figure 2.**
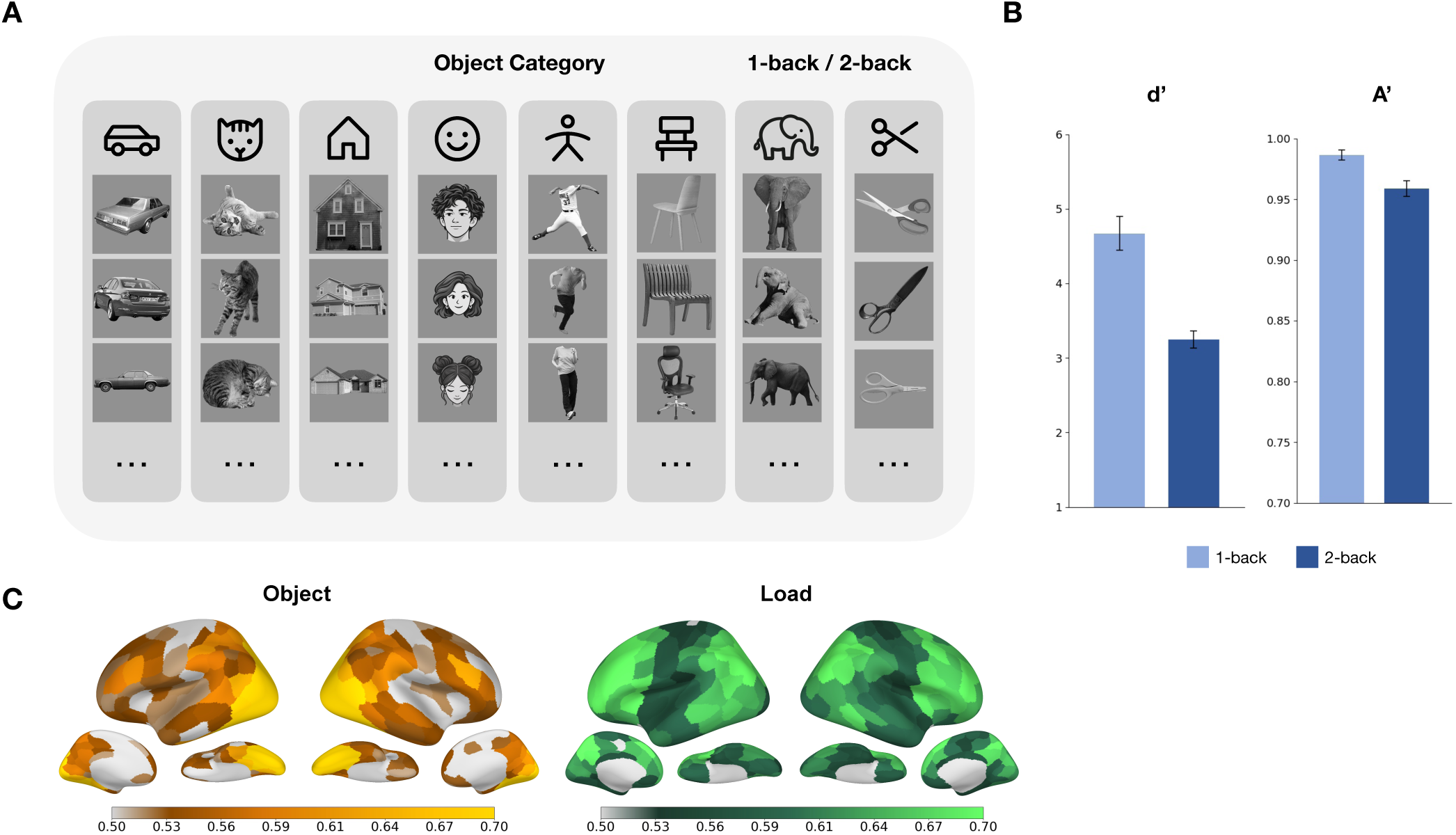
Pattern Decoding of Attention and Load in Experiment 2. **A:** The eight object categories from which trial stimuli were drawn. Example images are shown for each category. The cartoon faces shown here were generated using ChatGPT for illustration; the actual experiment used photographs of real human faces. Participants performed a standard 1-back and 2-back task. **B:** d’ and A’ behavioral performance under each of the two task conditions (n=12). **C**: Whole-brain maps showing cortical regions with significant average decoding of object category and load. Decoding accuracy in each region was tested against chance level (0.5), using one-sample t-tests followed by FDR correction across all ROIs in each hemisphere.

**Supplementary Figure 3.**
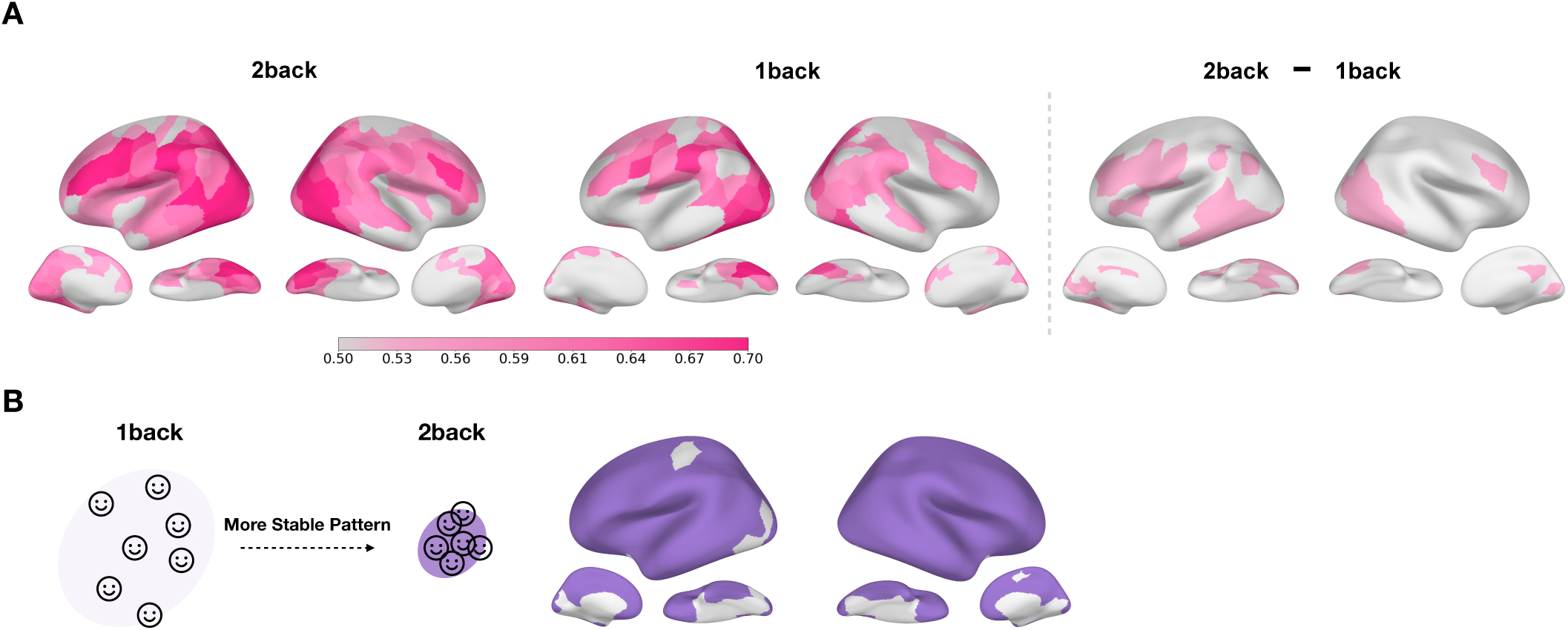
Load Effect Generalizes to the Representation of Attention. **A**: Whole-brain decoding maps for attention under different load conditions. Contrast map for attention decoding between loads is shown on the right. Decoding accuracy at each region was tested against chance level (0.5) using one-sample *t*-tests, followed by FDR correction for multiple comparisons across all ROIs in each hemisphere. Contrast maps were generated using one-sided t-tests followed by FDR correction across all ROIs in each hemisphere. Regions that did not survive correction are shown in light gray. **B**: Cortical regions showing more stable (less variable) attention representations from low to high load (p’s < 0.05, FDR-corrected across all ROIs in each hemisphere).

**Supplementary Figure 4.**
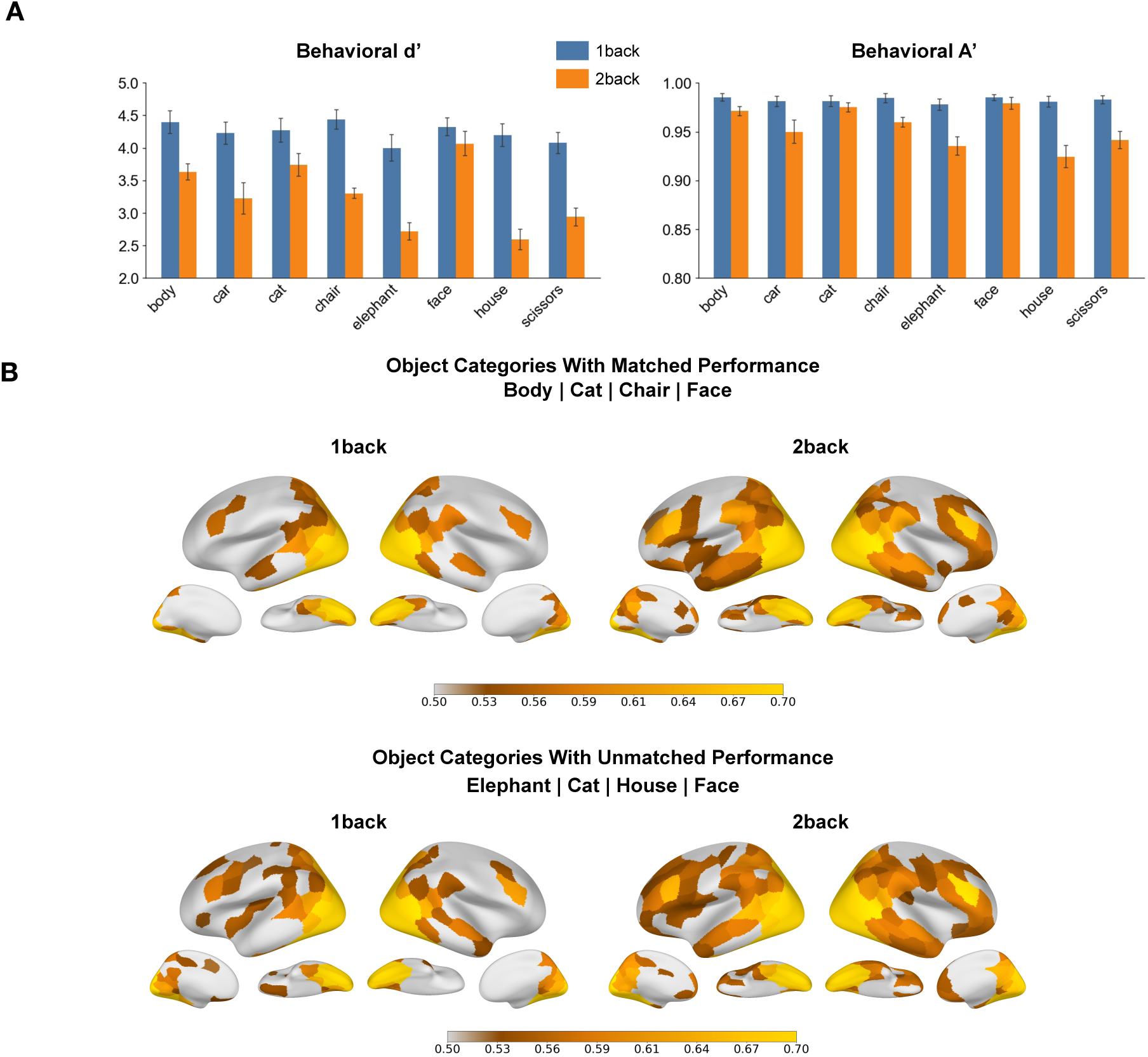
MVPA Decoding of Object Representations in Experiment 2, Controlling for Behavioral Performance. **A**: Average behavioral performance (d′ and A′) for each object category under each load condition. **B**: Whole-cortex object category decoding for subsets of object categories. **Top:** Decoding among the four more behaviorally matched categories (body, chair, cat, and face). **Bottom:** Decoding among the four more behaviorally unmatched categories (elephant, cat, house, and face). Colored region are those that exceed significance (*p* < 0.05, FDR-corrected across all regions in each hemisphere).

**Supplementary Figure 5.**
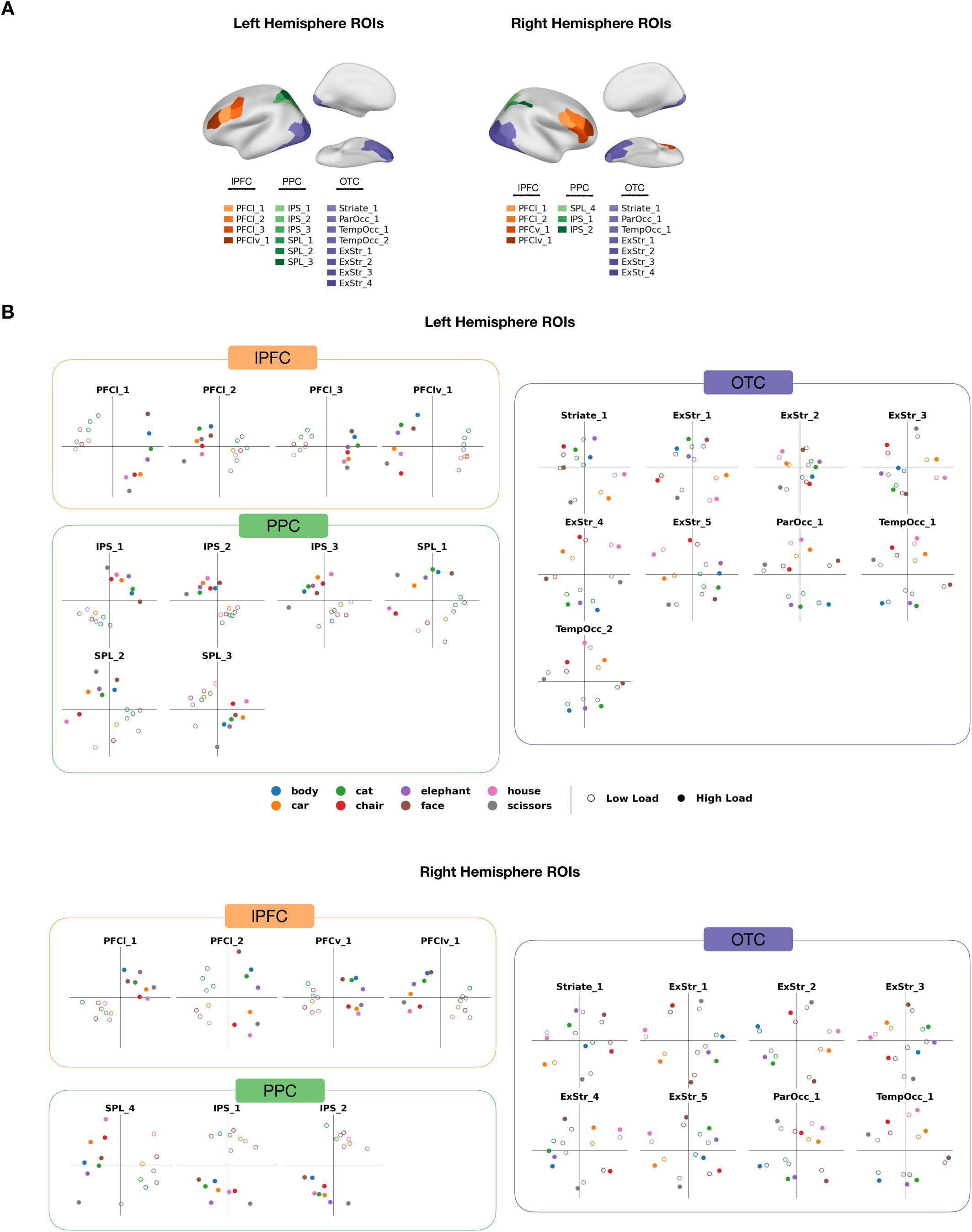
Representational Structure Across Selected Cortical ROIs in Experiment 2. **A**: Location of the selected cortical ROIs (based on Schaefer 200 parcellation) in the lPFC, PPC, and OTC for each hemisphere. **B**: MDS plots showing the representational structure of the 16 object-by-load conditions within selected left and right hemisphere ROIs. Dot color indicates object category, while marker fill (filled or hollow) indicates load condition.

**Supplementary Figure 6.**
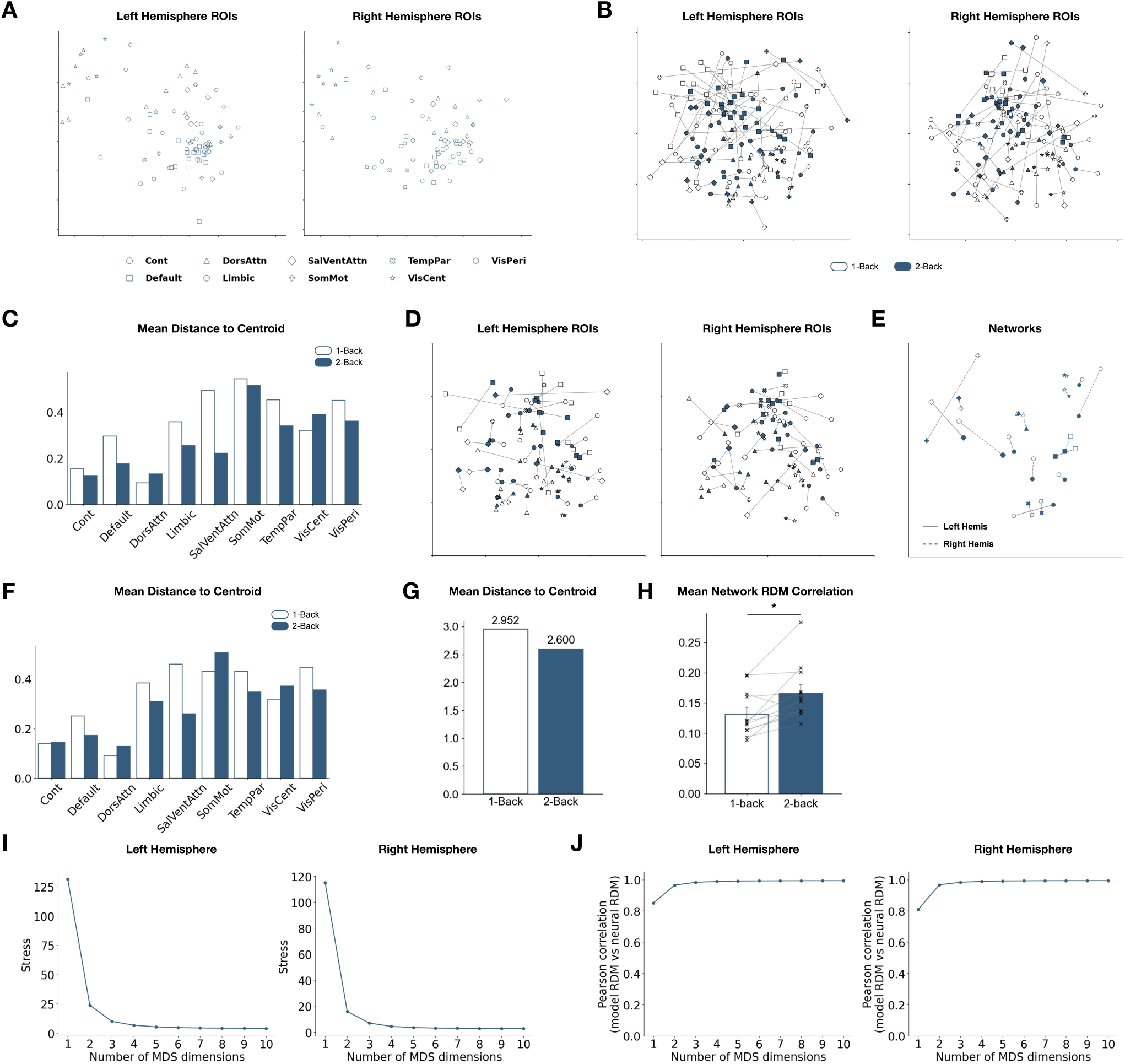
Higher Load Leads to a More Aligned Object Representational Space. **A:** MDS visualization of ROI-level RDMs. The 16 × 16 object-category RDMs from ROIs in both hemispheres were used to construct an ROI-level RDM, which was then projected into a common two-dimensional MDS space. The left and right hemispheres are plotted separately for visualization. Each marker represents one network. Only ROIs showing significant average object-category decoding accuracy are included (p’s < 0.05, FDR-corrected across ROIs in the same hemisphere). **B.** Load-dependent shifts in object representational spaces for individual ROIs. Lines connect the 1-back and 2-back MDS coordinates of each ROI, indicating the direction and magnitude of representational change with increasing load. ROIs are marked according to their canonical functional network. Only ROIs showing significant average object decoding across load conditions are included. **C**: Distance to the center of gravity for each network under each load condition, using regions that show significant average object decoding across conditions. To quantify alignment, we computed the Euclidean distance between each network’s MDS coordinate and the centroid (mean coordinate across all networks) within a given load condition. Here the plot displays mean distance to the centroid for each network under each load **D**: Same as in A, but with a more stringent ROI inclusion criterion — only regions showing significant object decoding in both load conditions are included. **E**: Network-level object representational space shifts, only regions showing significant object decoding in both load conditions are included. Solid and dashed lines indicate the left and right hemispheres, respectively. **(F-H)**: Distance to the center of gravity for each network under each load condition, using only regions that show significant object decoding in both load conditions. **F**: mean distance to the centroid for each network under each load. **G**: average distance across networks for each load. **H**: average RDM correlation across networks under each load (t = −3.406, p = 0.0059). **(I-J)**: Dimensionality analysis of RDM structure. **I**: stress plots (MDS fit) for each hemisphere’s ROI-by-ROI RDMs. **J**: correlation between the original ROI-by-ROI RDM and the reconstructed RDM as a function of the number of MDS components.

**Supplementary Figure 7.**
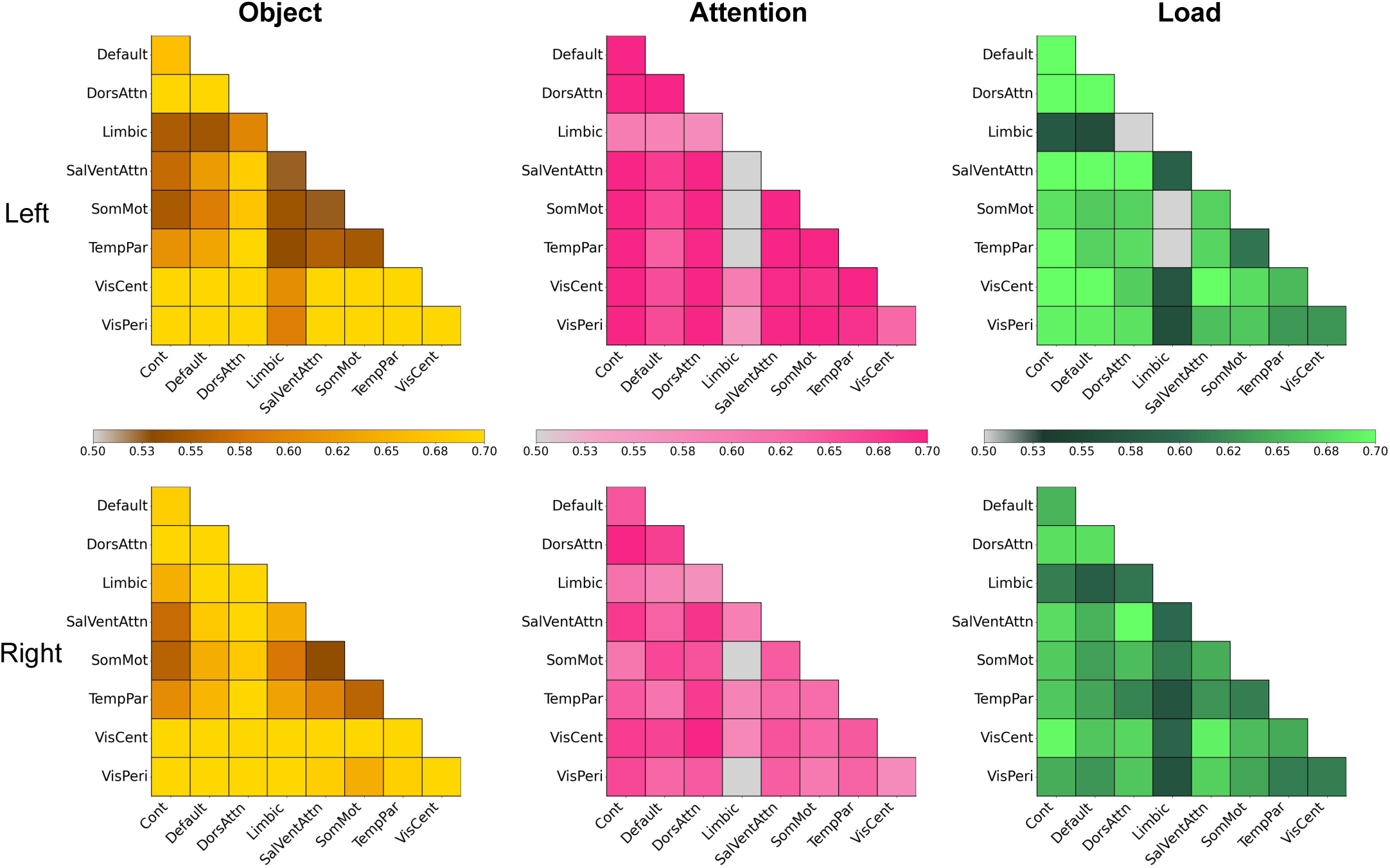
Decoding Object, Attention, and Load with Functional Connectivity Pattern Analysis (FCPA). FCPA decoding accuracy of object category (yellow), attention (pink), and load (green) between selected pairs of networks across the brain. Colored region pairs are those that exceed significance (*p* < 0.05, FDR-corrected across all pairs in each hemisphere).

**Supplementary Figure 8.**
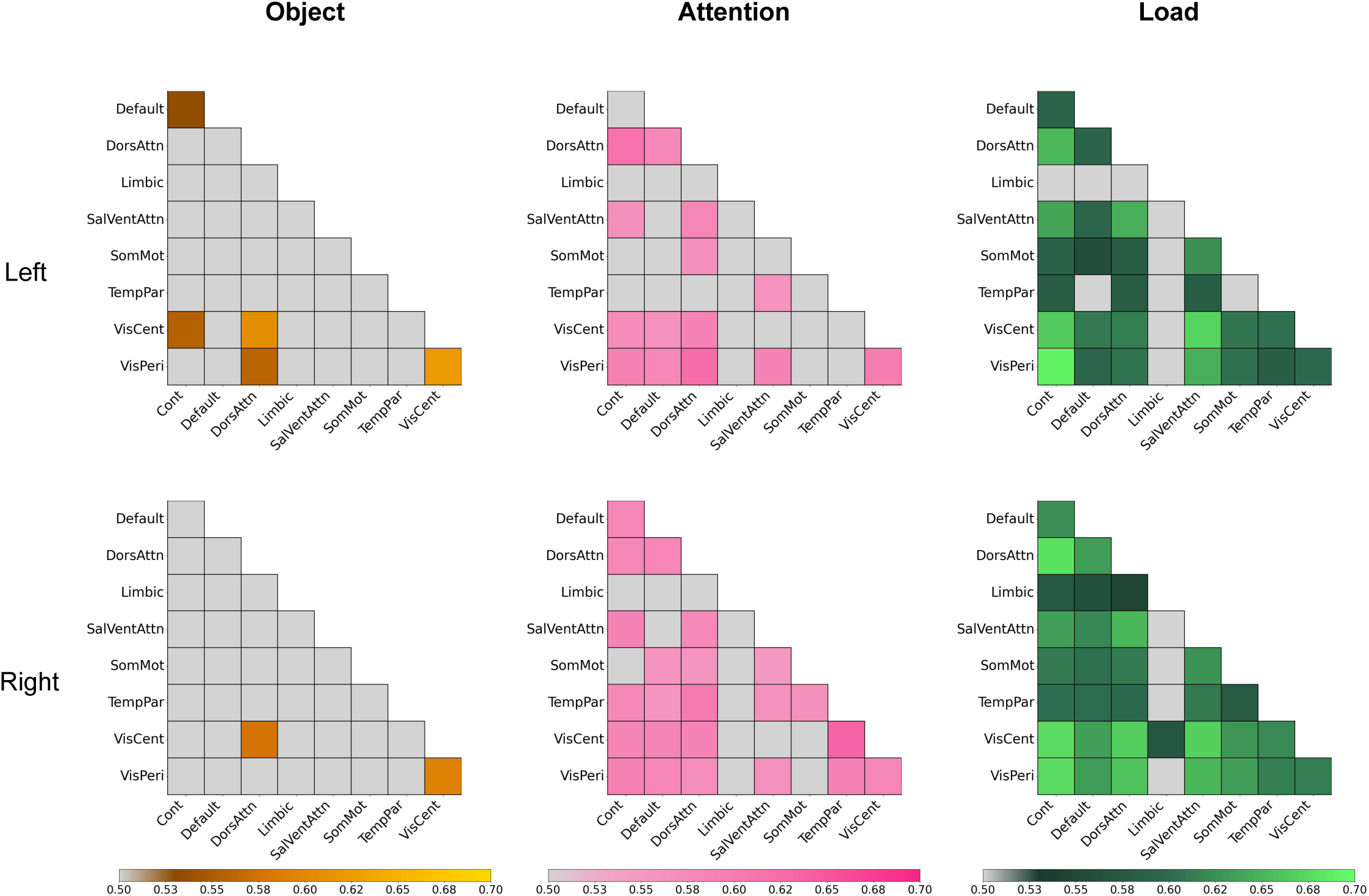
Decoding Object, Attention, and Load with Functional Connectivity Pattern Analysis (FCPA), Regressing Out Network Voxel Patterns with Bottom 500 Univariate Voxels. FCPA decoding accuracy after removing univariate and network multivariate voxel information for every pair of functional networks (top row) and right hemisphere (bottom row) ROIs. Univariate information was minimized by selecting the bottom 500 voxels that show least selectivity between conditions. Within-network multivariate patterns were regressed out from the network-pair functional connectivity patterns. Colored region pairs are those that exceed significance (*p* < 0.05, FDR-corrected across all pairs in each hemisphere).

**Supplementary Figure 9.**
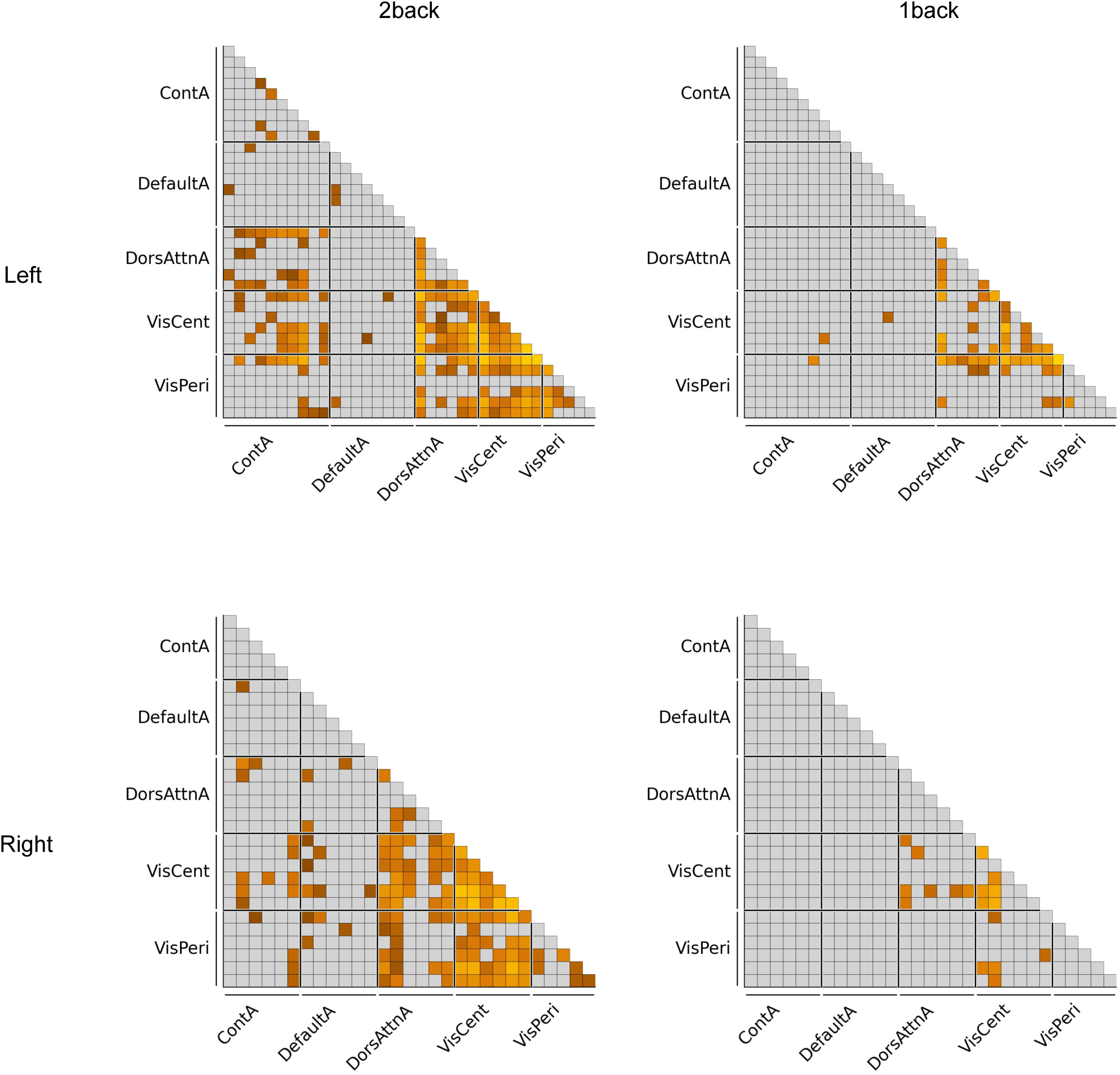
Decoding Object, Attention, and Load with Functional Connectivity Pattern Analysis (FCPA) Within Selected Network ROIs, Regressing Out Network Voxel Patterns Using the Bottom 500 Univariate Voxels. FCPA decoding accuracy after removing univariate and network multivariate voxel information for pairs of ROIs in a subset of selected networks (Control A, Default A, Dorsal Attention A, Visual Central, Visual Peripheral). Univariate information was minimized by selecting the bottom 500 voxels that show least selectivity between conditions. Within-network multivariate patterns were regressed out from the network-pair functional connectivity patterns. Colored region pairs are those that exceed significance (*p* < 0.05, FDR-corrected across all pairs in each hemisphere).

**Supplementary Figure 10.**
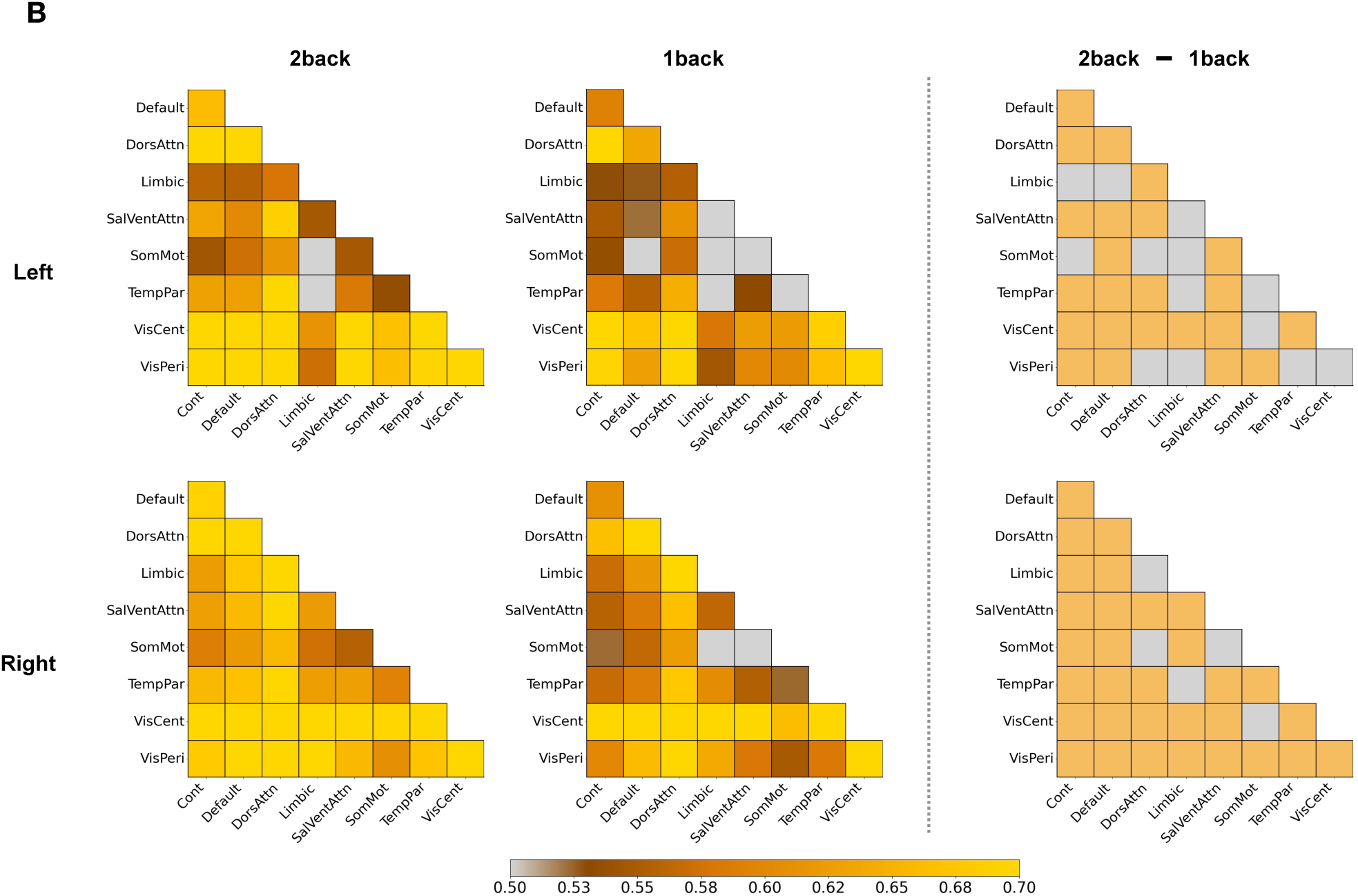
Decoding Object Category with Functional Connectivity Pattern Analysis (FCPA) Between Network Pairs Using Experiment 2 Data. Decoding accuracy for each network pair was tested against chance level (0.5) using one-sample t-tests, followed by FDR correction across all network pairs within the same hemisphere. Contrast maps were generated using one-sided t-tests, followed by FDR correction across the same set of network pairs. Network pairs that did not survive correction are shown in light gray.

**Supplementary Figure 11.**
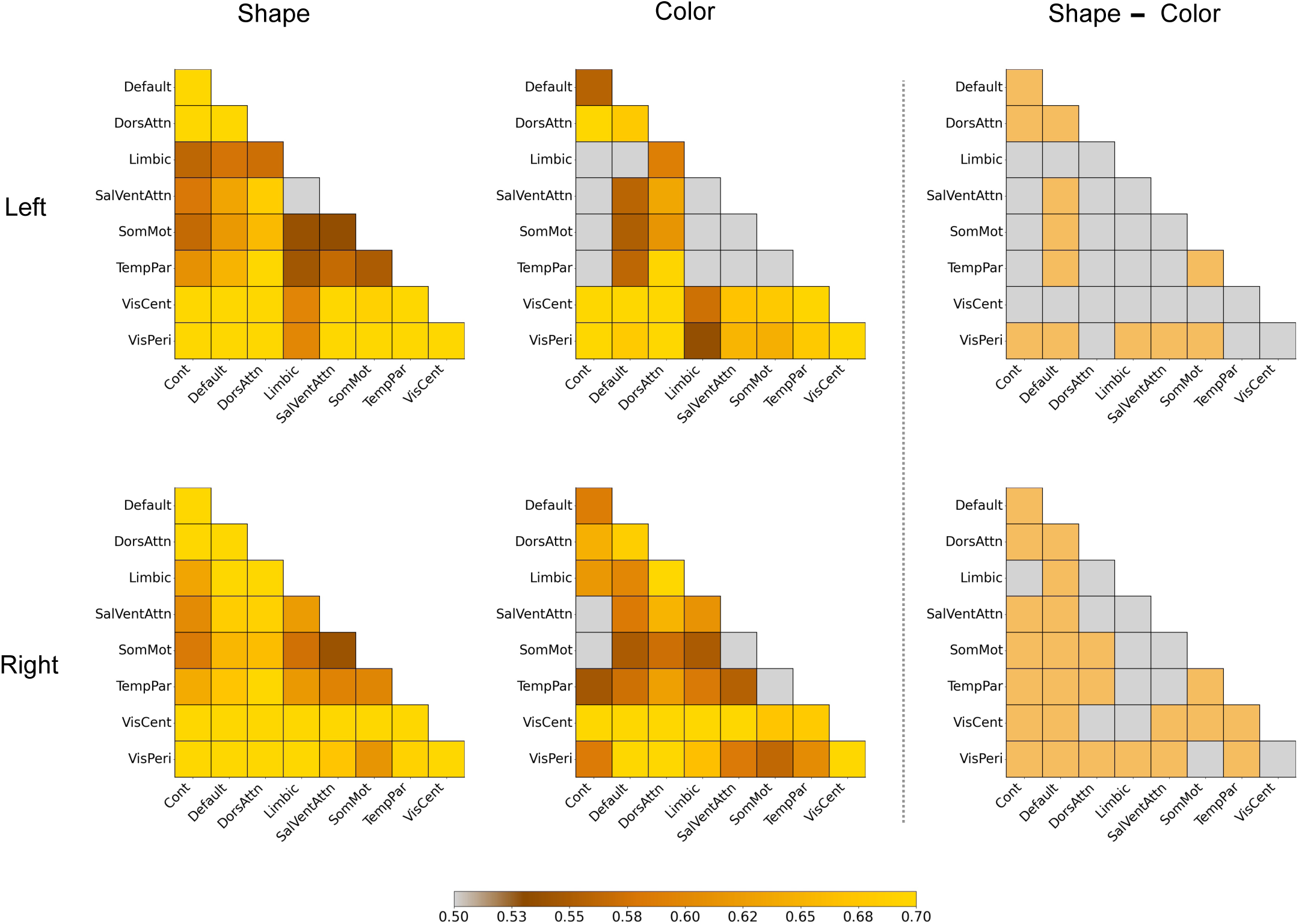
FCPA Decoding of Object Category Under Two Attention Conditions. **A**: FCPA decoding accuracy of object category under different attention conditions for left and right hemisphere networks. Decoding accuracy in each pair of networks was tested against chance level, 0.5, using one-sample t-tests followed by FDR correction for multiple comparisons. Contrast maps were generated using one-sided t-tests followed by FDR correction across ROIs in each hemisphere. Regions that did not survive correction are shown in light gray.

**Supplementary Figure 12.**
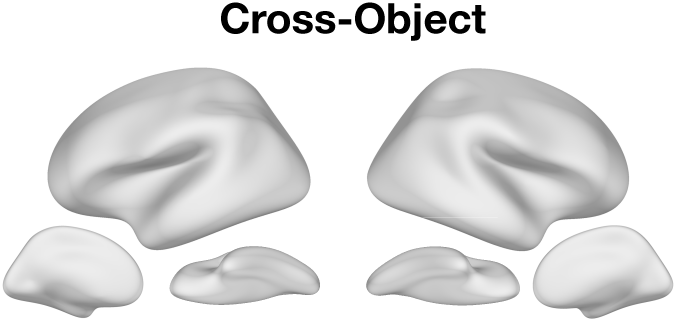
Load Decoding Across Object Categories, Experiment 2. Binary decoding drop maps for load across object categories, shown separately for each load condition. For each ROI, we conducted paired t-tests comparing attention decoding accuracy within the same object category to decoding across different object categories. No ROI exhibits significantly reduced decoding performance under cross-object decoding (p < 0.05, FDR-corrected across all ROIs in each hemisphere), indicating object-independent load representations.

**Supplementary Figure 13.**
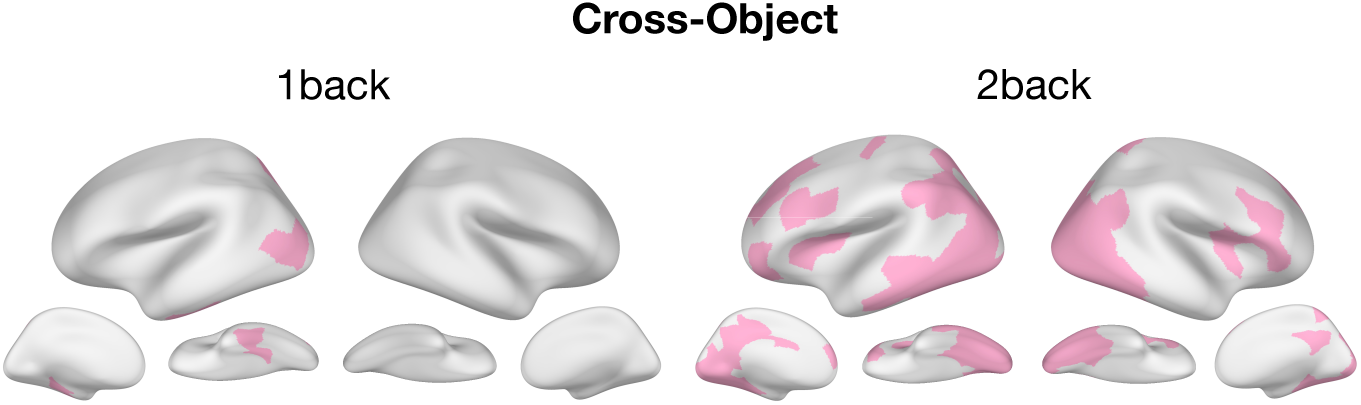
Attention Decoding Across Object Categories. Binary decoding drop maps for attention across object categories, shown separately for each load condition. For each ROI, we conducted paired t-tests comparing attention decoding accuracy within the same object category to decoding across different object categories. ROIs shown in pink exhibit significantly reduced decoding performance under cross-object decoding (p < 0.05, FDR-corrected across all ROIs in each hemisphere), indicating object-dependent attention representations. Regions not surviving correction are shown in light gray.

**Supplementary Figure 14.**
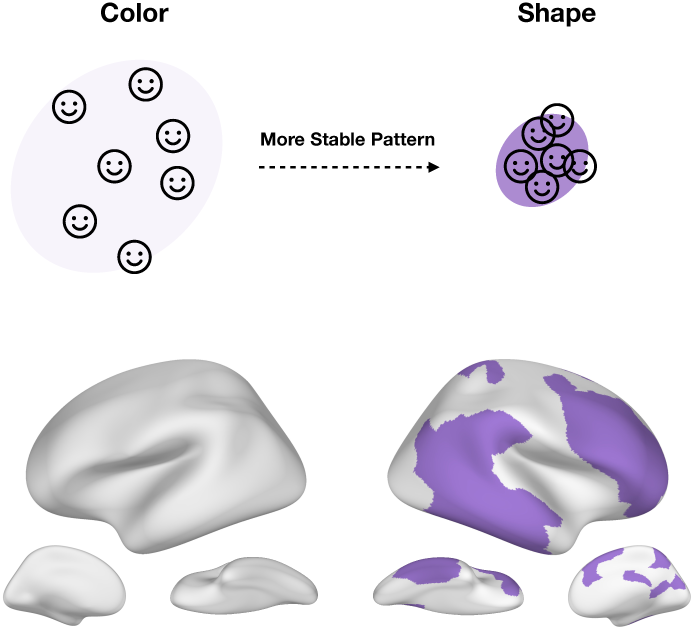
Load Pattern Stability Analysis for Experiment 1. Purple areas correspond to cortical regions exhibiting more stable (less variable) load representations across attention conditions (color vs. shape; p’s < 0.05, FDR-corrected across all ROIs in each hemisphere).

